# Immunosuppression and Outcomes in Acute Myeloid Leukemia

**DOI:** 10.1101/2021.09.03.458879

**Authors:** F Ferraro, CA Miller, KA Christensen, NM Helton, M O’Laughlin, CC Fronick, RS Fulton, J Kohlschmidt, AK Eisfeld, CD Bloomfield, SM Ramakrishnan, RB Day, LD Wartman, GL Uy, JS Welch, MJ Christopher, SE Heath, JD Baty, MJ Schuelke, JE Payton, DH Spencer, MP Rettig, DC Link, MJ Walter, P Westervelt, JF DiPersio, TJ Ley

## Abstract

Acute myeloid leukemia (AML) patients rarely have long first remissions (> 5 years) after standard-of-care chemotherapy, unless classified as favorable risk at presentation. Identification of the mechanisms responsible for long vs. more typical, short remissions may help to define prognostic determinants for chemotherapy responses. Using exome sequencing, RNA-sequencing and functional immunologic studies, we characterized 28 Normal Karyotype (NK)-AML patients with >5 year first remissions after chemotherapy (Long First Remissions, LFR) and compared them to a well-matched group of 31 NK-AML patients who relapsed within 2 years (Standard First Remissions, SFR). Our combined analyses indicated that genetic risk profiling at presentation (as defined by ELN 2017 Criteria) was not sufficient to explain the outcomes of many SFR cases. Single cell RNA-sequencing studies of 15 AML samples showed that SFR AML cells differentially expressed many genes associated with immune suppression. The bone marrow of SFR cases had significantly fewer CD4+ Th1 cells; these T-cells expressed an exhaustion signature and were resistant to activation by T-cell receptor stimulation in the presence of autologous AML cells. T-cell activation could be restored by removing the AML cells, or blocking the inhibitory MHC Class II receptor, LAG3. Most LFR cases did not display these features, suggesting that their AML cells were not as immunosuppressive. These findings were confirmed and extended in an independent set of 50 AML cases representing all ELN 2017 risk groups. AML cell-mediated suppression of CD4+ T-cell activation at presentation is strongly associated with unfavorable outcomes in AML patients treated with standard chemotherapy.

## Introduction

A normal karyotype is the most common cytogenetic finding in AML cells at presentation (“NK-AML”), and has long been associated with an intermediate risk for relapse (1). Although most NK-AML patients achieve a complete remission following standard induction chemotherapy (2, 3), the majority of patients will experience relapse within 2 years, and fewer than 10% will remain in remission beyond 5 years without an allogeneic transplant (4). Refinements introduced with the ELN 2017 Criteria (presence or absence of mutations in *NPM1*, *FLT3*-ITD, *ASXL1*, *RUNX1*, *TP53* and/or biallelic *CEBPA* mutations) have helped to reclassify some NK-AML patients into the favorable or adverse risk group categories. However, the mutational heterogeneity of NK-AML, combined with our limited understanding of the biological consequences deriving from the interplay among mutations, still poses a major challenge for risk stratification at presentation, and for post-remission treatment decisions (5, 6).

The genetic and epigenetic characteristics of NK-AML patients with very long first remissions have not yet been defined, and it is not clear whether these patients are living free from disease, or whether they harbor “dormant” AML cells that may be held in check by immune surveillance or cell-intrinsic mechanisms. Predictive algorithms for identifying these very long responders at presentation do not yet exist, but clearance of all AML-associated mutations assessed after induction chemotherapy is strongly associated with prolonged relapse-free and overall survival (7–10). Persistent ancestral clones have been detected in patients in complete morphologic remission within the first 2-3 years after treatment (11, 12), but similar studies of patients with very long first remissions have not yet been described.

Most published AML studies have justifiably focused on the mechanisms responsible for treatment failure. However, the reasons for chemotherapy successes remain underexplored, even though they hold the potential to uncover determinants that are especially important for excellent outcomes--and that might be identifiable at diagnosis. To address this, we have carefully studied two well-matched groups of NK-AML patients that were selected retrospectively for dramatically different outcomes after chemotherapy (long vs. standard first remissions). In this study, we show that the outcomes of AML patients are not only related to their mutational status, but also to an immunosuppressive phenotype of AML cells at presentation (or lack thereof), a finding that was predicted 50 years ago by Freireich and colleagues (13), and extended in this report.

## Methods

### Patient Selection

All clinical data are available in the **Supplementary Appendix, Sections A and B**. NK-AML patients were selected for this study because their outcomes are highly variable and still difficult to predict at presentation. We defined Long First Remissions (LFRs) as lasting more than 5 years to ensure the durability of treatment responses, which increases the chance of identifying robust biomarkers associated with excellent responses. The comparator set of Standard First Remissions (SFRs) were defined as having an initial relapse within 2 years of presentation, since 70% of NK-AML cases relapse within 2 years. We did not use the ELN 2017 Criteria to select the initial sample sets for the study, since many of the patients were treated before gene testing for common AML mutations was routinely performed, and it did not guide therapeutic decisions. We required the following inclusion criteria for all patients: 1) morphologically documented *de novo* AML with adequate bone marrow and matched control (skin) DNA for sequencing studies, 2) at least 18 years of age, 3) normal karyotype of AML cells at presentation, 4) received “7+3” (or similar variants) for induction chemotherapy, and 5) received high or intermediate-dose AraC as the primary consolidation therapy (i.e. no allogeneic transplant in first remission). For the LFR cases, the initial remission had to last at least 5 years. For the SFR cases, patients either had a morphologically documented relapse occurring within 2 years of starting therapy (28 patients) or primary refractory disease with 7+3 (3 patients), prior to any form of transplant (none of these patients were transplanted in first remission). To identify the LFR samples for study, 1,579 cases were screened from our AML collection at Washington University, and 19 cases meeting all criteria were identified (1.2%). 9 additional LFR cases matching the above criteria were obtained after screening 846 NK-AML samples (1.06%) at the Alliance collection housed at The Ohio State University (but matched normal DNA and RNA was not available for these samples). Samples were acquired as part of studies that were approved by the Human Research Protection Office at both institutions. Thirty one well-matched NK-AML cases with SFRs were selected as comparators. Post-hoc analyses confirmed that there were no significant differences in age, sex, bone marrow blast percentage at diagnosis, karyotype, or induction/consolidation treatments between LFR and SFR patients. All the patients provided written informed consent that included explicit permission for genetic studies, under IRB-approved protocols (# 201011766 for Washington University and CALGB / Alliance 9665 [NCT0089922]) and 20202 [NCT00900224] for samples obtained from The Ohio State University).

### Survival analysis

Overall survival (OS) was defined as the time from diagnosis to death from leukemia. Patients who died in remission were censored at the time of death. Relapse free survival (RFS) was defined as the time from diagnosis to relapse or death from leukemia, whichever occurred first. Patients who were alive and disease-free were censored at last follow-up. Thirteen out of 50 cases in the extension cohort who were transplanted in first remission were censored at the time of transplant. The distributions of OS or RFS between different groups were described using Kaplan-Meier product limit methods, and compared by a log-rank test. Survival curves were visualized with GraphPad prism (v. 9.0.2).

### Molecular, Immunologic, and Statistical Analyses

DNA, RNA, and viably cryopreserved cells were obtained from the bone marrow aspirates of normal donors or AML patients at presentation, or at various time points during remission. The LFR and SFR samples that were used for sequencing and functional studies are listed in the Supplementary Appendix. For full details regarding T-cell activation studies, flow cytometry, cell culture, sequencing, bioinformatic and statistical analyses, refer to the Supplementary Appendix.

## Results

### Mutational spectrum of NK-AML cases with Long vs. Standard First Remissions

The clinical characteristics of the 28 LFR and 31 SFR cases are summarized in **Table 1**, and detailed in **Supplementary Table 1**. All patients had a normal karyotype defined by standard cytogenetics, and received standard-of-care induction chemotherapy and consolidation, as detailed in the Patient Selection section. At the time of this writing, none of the LFR patients from Washington University had relapsed (range 5.4-17+ years, median follow up 8.7 years). The two groups were comparable in terms of age, sex, percentage of bone marrow blasts and white blood cell counts at diagnosis (**Table 1**). The Kaplan-Meier (KM) distribution of relapse free survival (RFS) and overall survival (OS) of the LFR and SFR cases is shown in **Figures 1A** and **B**). The mutational landscape of these cases was defined by sequencing the exomes from the presentation bone marrow samples from all cases^5^. Matched normal (skin) samples were available for the 19 LFR and 31 SFR cases banked at Washington University. Somatic mutation status was inferred for the 9 Alliance samples by limiting mutation calling to known, recurrently mutated AML genes^14^ (**Figure 1C, Supplementary Table 2)**.

**Table 1.** Clinical characteristic of the Standard versus Long First Remission patients

|  |  |  |  |
| --- | --- | --- | --- |
|  | **SFR (n=31)** | **LFR (n=28)** | **p value** |
| **Age at diagnosis - yr** |  |  | 0.93 |
| Median | 51 | 55 |  |
| Range | 21-76 | 19-71 |  |
| **Female sex (%)** | 48.40 | 42.90 | 0.79 |
| **WBC count** |  |  | 0.09 |
| Median | 52.9 | 26.8 |  |
| Range | 1.5-222.8 | 1.6-298.4 |  |
| **Blasts in the BM (%)** |  |  | 0.28 |
| Median | 72 | 72 |  |
| Range | 30-97 | 23-91 |  |
| **Relapse Free Survival- months** |  |  | <0.001 |
| Median | 7.7 | 104.1 |  |
| Range | 1.1-20.6 | 65.1-199.1 |  |
| **Overall Survival - months** |  |  | <0.001 |
| Median | 17 | 104.1 |  |
| Range | 2.2-188.1 | 65.1-199.1 |  |
| **ELN category** |  |  |  |
| Favorable | 14 (45%) | 26 (93%) | <0.001 |
| Intermediate | 15 (48%) | 1 (3.5%) | <0.001 |
| Adverse | 2 (8%) | 1 (3.5%) | 1 |
| **Occurrence of mutation (%)** |  |  |  |
| *NPM1* | 17 (54.8%) | 24 (85.7%) | 0.01 |
| *DNMT3A* | 14 (45.2%) | 11 (39.3%) | 0.79 |
| *FLT3-ITD* | 14 (45.2%) | 5 (17.9%) | 0.03 |
| *other FLT3* | 5 (16.1%) | 6 (21.4%) | 0.74 |
| *MYC* | 1 (3.2%) | 6 (21.4%) | 0.05 |
| *NRAS* | 5 (16.1%) | 11 (39.3%) | 0.08 |
| *WT1* | 6 (19.4%) | 3 (10.7%) | 0.48 |
| *TET2* | 4 (12.9%) | 3 (10.7%) | 1 |
| *PTPN11* | 2 (6.5%) | 5 (17.9%) | 0.24 |
| *IDH1* | 4 (12.9%) | 1 (3.6%) | 0.36 |
| *IDH2* | 3 (9.7%) | 3 (10.7%) | 1 |
| *CEBPA* | 4 (12.9%) | 3 (10.7%) | 1 |

**Figure 1.**
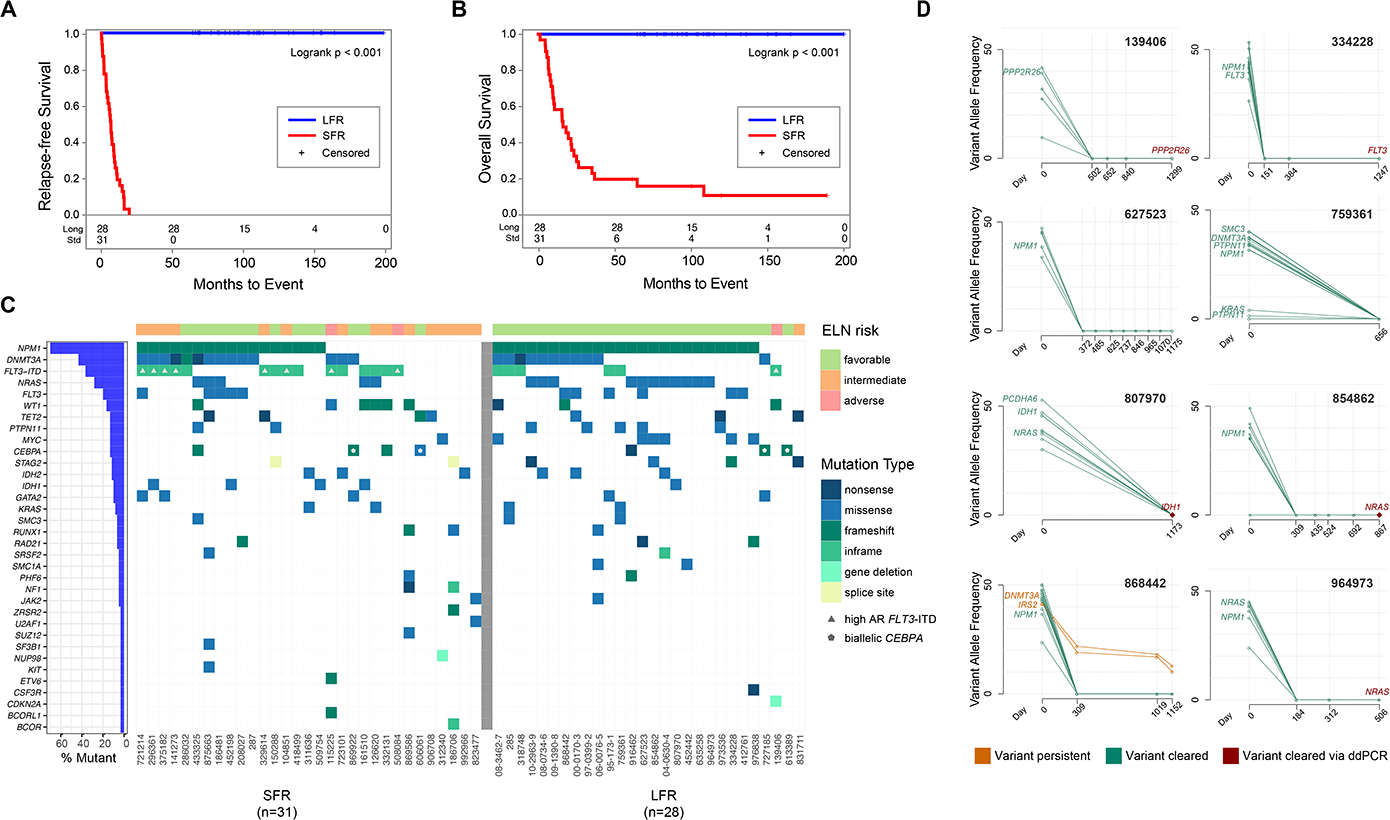
Clinical and genomic features of AML patients at presentation and in remission. **Panel A,** relapse-free survival and **Panel B,** overall survival curves for NK-AML patients who were treated with chemotherapy only for induction and consolidation. The blue line represents the long first remission cases (LFR, n=28) and the red line the standard first remission cases (SFR, n=31). **Panel C** shows the mutational landscape and the ELN classification for each case. Each column represents a patient, and each row represents a gene that is mutated in at least one of these cases. Every case had one or more recognized AML driver mutations, with a median of 11 (range 1-37) protein-altering somatic mutations per case in the Washington University samples. Color indicates the type of mutation, as specified in the legend. Cases with high *FLT3*-ITD allelic ratio are indicated by the grey triangle in the figure. Blue bars at left indicate the mutation frequency in the sample set. **Panel D** shows clearance plots displaying the Variant Allele Frequencies (VAFs) of recurrently mutated AML genes, plotted at presentation (day 0) and at available time points during clinically defined remissions (days), assessed with error-corrected sequencing. The average coverage of all variants for the remission samples was 3042x, with a range of 161-13,266x. Mutation clearance of the genes highlighted in red was confirmed with digital droplet PCR, at a sensitivity of ~1 AML per 100,000 cells tested (**Supplementary Figure 2**).

Using these data, we classified patients according to the ELN 2017 genetic risk categories (**Table1, Supplementary Table 1 and Figure 1C**). The ELN Criteria classified 26 of the 28 (93%) LFR cases as favorable risk, 1/28 (3.5%) as intermediate risk and 1/28 (3.5%) as adverse risk. For the 31 SFR cases, 14/31 cases (45%) were classified as favorable risk (median RFS 301 days), 15 (48%) were classified as intermediate risk (median RFS 195 days), and 2 cases (8%) were classified as adverse risk by ELN criteria (median RFS 330 days). Kaplan-Meyer plots for the ELN-classified SFR cases are shown in **Supplementary Figure 1**. The fact that nearly half of the SFR patients with short remissions are classified as favorable risk according to ELN criteria suggested that other factors must play a role in the fates of the SFR patients treated with standard of care chemotherapy alone.

### Evaluation of Persistent Molecular Disease in Long First Remission cases

For eight of the LFR cases, we had access to remission samples obtained 506-1299 days after presentation. We performed error-corrected sequencing (sensitivity of 4 AML cells in 10,000 total cells) of the presentation and remission samples, targeting 5 to 18 known somatic variants per case(14) (**Supplementary Table 3**). For 7 of the 8 cases, no AML-associated mutations could be detected in any remission sample (**Figure 1D**). In one case (868442), a *DNMT3A*^R882H^ and an *IRS2*^D106Y^ mutation persisted in remission, suggesting that the patient retained a pre-leukemic, ancestral clone. To increase the detection sensitivity of persistent AML cells to ~1 in 100,000 (0.001%), we performed digital droplet PCR for a single founding clone mutation (for which commercial reagents were available) for 5 patients in remission; no persistent AML cells bearing these mutations was detected in any of these samples (**Supplementary Figure 2**).

### RNA-sequencing analyses of Standard vs. Long First Remission cases

Because bulk RNA sequencing cannot accurately define the expression patterns of cellular subsets in AML samples, we performed single-cell RNA-sequencing (scRNA-seq) of unfractionated bone marrow from the presentation samples of 8 LFR and 7 SFR cases that were chosen based on high cellular viability (>50%) in available cryovials; the two sets had comparable fractions of myeloblasts based on bone marrow aspirates flow cytometry at diagnosis (mean for LFR 71.1%, vs. SFR 66.3%, p=0.51). In addition, whole bone marrow and CD34+ flow-sorted cells were evaluated from 2 healthy donors (ages 33 and 36). **Supplementary Figures 3A and 3B** display UMAP projections of normalized expression data from all cells, labeled by cell type (15) and sample, respectively. Non-malignant cells, such as immune cells and erythroid cells, formed common clusters containing cells from all 17 samples. For each AML sample, we identified cells expressing known AML-associated somatic mutations (defined by prior DNA sequencing), allowing us to restrict our analysis to genetically-defined AML cells (**Figure 2A**) (16). Targeted investigation of cell type proportions (**Figure 2B**) and pathways expected to be relevant for chemotherapy sensitivity (i.e. cell cycle, DNA repair, or AraC responsiveness) revealed no significant differences between LFR and SFR AML cells (**Supplementary Figures 3C through F, Supplementary Table 4)**.

**Figure 2.**
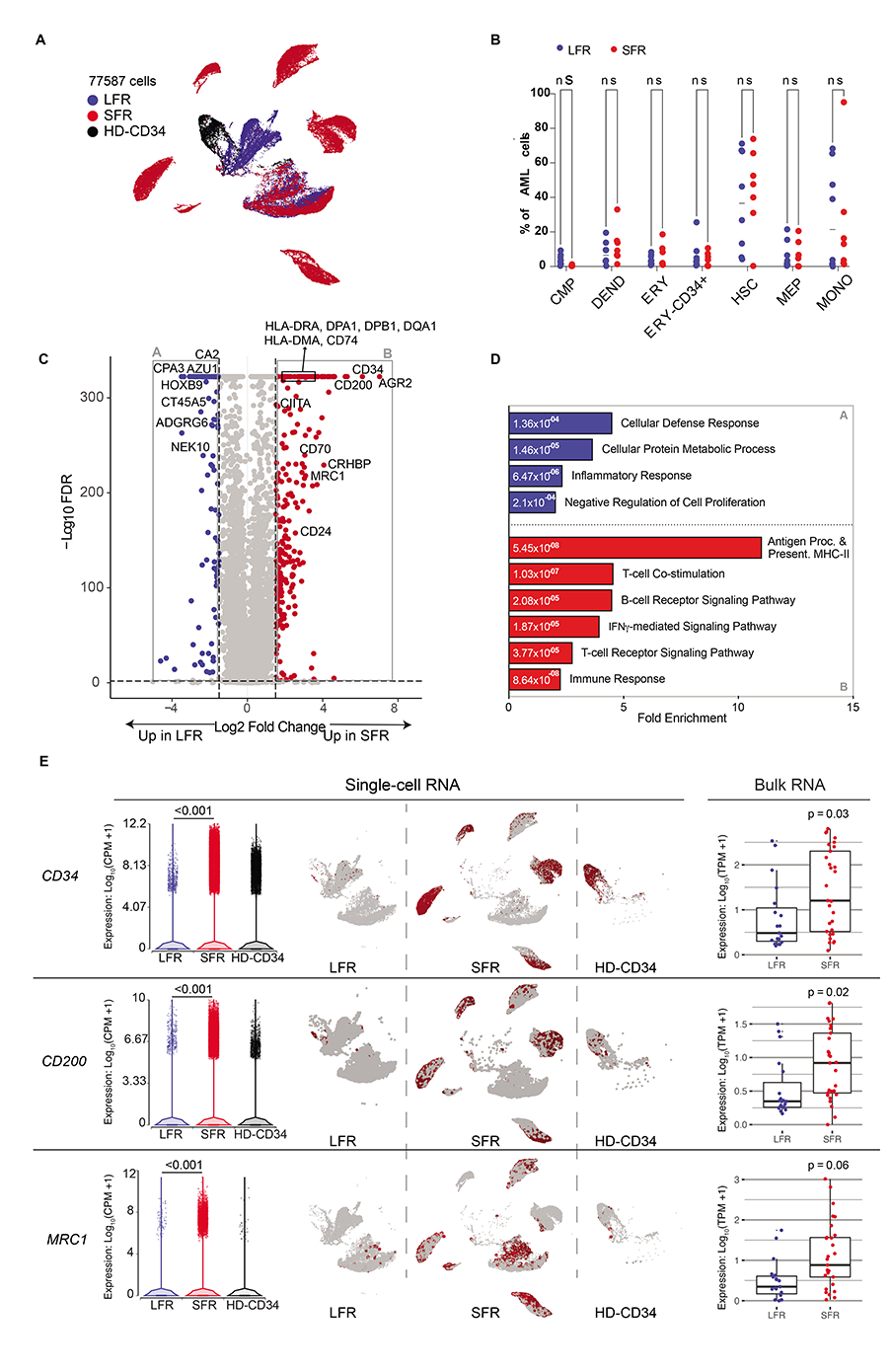
RNA-sequencing studies of LFR and SFR AML samples. **Panel A**: UMAP projection of 77,587 genetically-defined AML cells from 8 LFR and 7 SFR samples at presentation, and purified CD34+ cells from the bone marrow samples of 2 healthy donors (HD-CD34). Colors reflect the sample groups, as indicated in the legend. **Panel B:** Proportion of the genetically-defined AML cells composed of different cell types across each AML sample for LFR cases (blue) vs. SFR cases (red). Horizontal lines indicate median values. **Panel C**: Differentially Expressed Genes (DEGs) in genetically-defined AML cells from the presentation bone marrow samples of SFR vs. LFR patients. Dashed lines indicate log2 fold change of ± 2 and FDR<0.001. Red points represent genes with significantly higher expression in the SFR cases, and blue points represent genes that are more highly expressed in the LFR cases. **Panel D:** Enrichment for Gene Ontology terms in the DEGs. Blue and red bars are pathways enriched in the LFR or SFR cases, respectively; numeric values indicate FDR. **Panel E**: Left: expression of *CD34*, *CD200*, and *MRC1* in single-cell data, with each point representing a cell. Center: UMAP plots (split by category) showing single-cell expression of the corresponding gene. Right: expression from bulk RNA-seq datasets with additional AML samples (LFR, blue dots, n=19; SFR, red dots, n=31).

Applying ANOVA to genetically-defined AML cells from LFR vs. SFR cases, we identified 11,657 differentially expressed genes (DEGs) (FDR=0.001, log FC>±1, **Figure 2C and Supplementary Table 5**). Functional enrichment analysis showed that morphogenesis, chemotaxis, and inflammation pathways were enriched in the genes that were more highly expressed in the LFR AML cells, while genes involved in MHC Class II-mediated antigen presentation, TCR signaling, and immune responses were expressed at significantly higher levels in the AML cells from SFR samples (**Supplementary Figure 4A and B**). Because *NPM1* mutations are known to influence MHC Class II (17) and HOX gene expression (18, 19), we repeated the ANOVA comparison after restriction to the *NPM1* mutant cases (n=8 LFR cases, and n=3 SFR cases, **Supplementary Table 6**). We identified 8,191 DEGs that were shared with the previous analysis (FDR=0.001, log FC±1, **Supplementary Figure 4C and Supplementary Table 6**). Repeat Gene Ontology analysis with the overlapping DEGs eliminated the HOX gene pathways as significant, but increased the significance of the MHC class II signature found in the SFR AML cells (**Supplementary Table 6 and Figure 2E**). To extend these findings, we evaluated MHC class II gene expression using the bulk RNA-seq data from the 31 SFR vs 19 LFR cases, and found that *CIITA* (the master regulator of MHC class II expression) and several Class II genes were expressed at significantly higher levels in the SFR cases (**Supplementary Figure 4D**). Although *NPM1* mutations are known to be associated with low CD34 expression (20–23), we observed that the LFR AML cells expressed the *CD34* gene at significantly lower levels than SFR AML cells, even when limiting the comparison to the *NPM1* mutated cases (**Figure 2E and Supplementary Table 7)**. Two immunomodulatory genes were found to be expressed at significantly higher levels in SFR AML cells: *CD200* and *MRC1* (**Figure 2E**). CD200 has been reported to be associated with immunosuppression and worse outcomes in AML (24–26). *MRC1* encodes CD206, a marker of tumor-promoting macrophages in chronic inflammatory diseases (27, 28) and solid tumors (29, 30); it was recently identified as an independent negative prognostic indicator in AML (31). The expression differences for *CD34*, *CD200*, and *MRC1* were similar in the bulk RNA sequencing data from the extended sample set (**Figure 2E**).

### Expression signatures in marrow-infiltrating immune cells

We next sought to compare the expression signatures of the infiltrating immune cells in the leukemic marrow samples of the LFR vs. SFR cases. B cells, Natural Killer cells, Regulatory T-cells, and gamma delta T-cell populations were too small to allow for meaningful studies of differential gene expression. However, we identified 9,004 T-cells after pooling all samples, shown in the UMAP projection of **Figure 3A**. To identify and label specific T-cell subpopulations, we applied a graph-based clustering approach (**Figure 3B, Supplementary Figure 5A and Supplementary Table 8**). SFR marrows at presentation had a smaller proportion of total T-cells (**Supplementary Figure 5B**). Flow cytometric evaluation of 15 LFR, 26 SFR, and 8 healthy adult donor marrow samples (age range 25-40) confirmed that the fraction of CD3+ T-cells in SFR AML bone marrows was significantly lower than that of LFR and healthy donor marrows (**Figure 3C**); specifically, SFR bone marrow samples contained a significantly smaller fraction of CD4+ T-cells (**Figure 3D**). Among the CD4 T-cell subsets, the SFR marrow samples had significantly fewer Th1 cells, but no statistical differences were detected in the proportions of Th2, Th17, or Treg cells, which were all rare populations in these AML samples (**Figure 3E**). We next tested the differential expression of the genes encoding 143 previously described T-cell specific activation/exhaustion markers (32, 33) in the total immune cells of SFR vs LFR samples. Notably, 57 out of the 62 differentially expressed genes were expressed at significantly higher levels in the SFR T-cells (**Figure 3F, and Supplementary Table 9**). These included genes encoding several putative immune checkpoint proteins, including *LAG3,* an inhibitory receptor for MHC Class II (log FC = 1.59, FDR<0.001), *CD38* (log FC = 3.28, FDR<0.001), and *HAVCR2* (which encodes TIM3; log FC = 1.51, FDR<0.01), although the latter was expressed by only a very small number of T-cells (**Supplementary Figure 5C**). The degree of differential expression for these genes varied across T-cell subpopulations; for both *LAG3* and *CD38* it was more prominent in the CD4+ T-cell compartment (**Figure 3G;** *LAG3,* log FC = +2.06, FDR<0.001; C*D38* log FC = +4.58, FDR<0.001). CD4+ T-cells did not express significant levels of *HAVCR2* (**Supplementary Figure 5D and Supplementary Table 10**). Other genes encoding potentially relevant immune checkpoint proteins, such as *PDCD1*, *TIGIT*, *CTLA4,* and *CD200R1,* were not differentially expressed by the immune cells of LFR vs. SFR samples (**Supplementary Figures 5D, Supplementary Table 9**).

**Figure 3.**
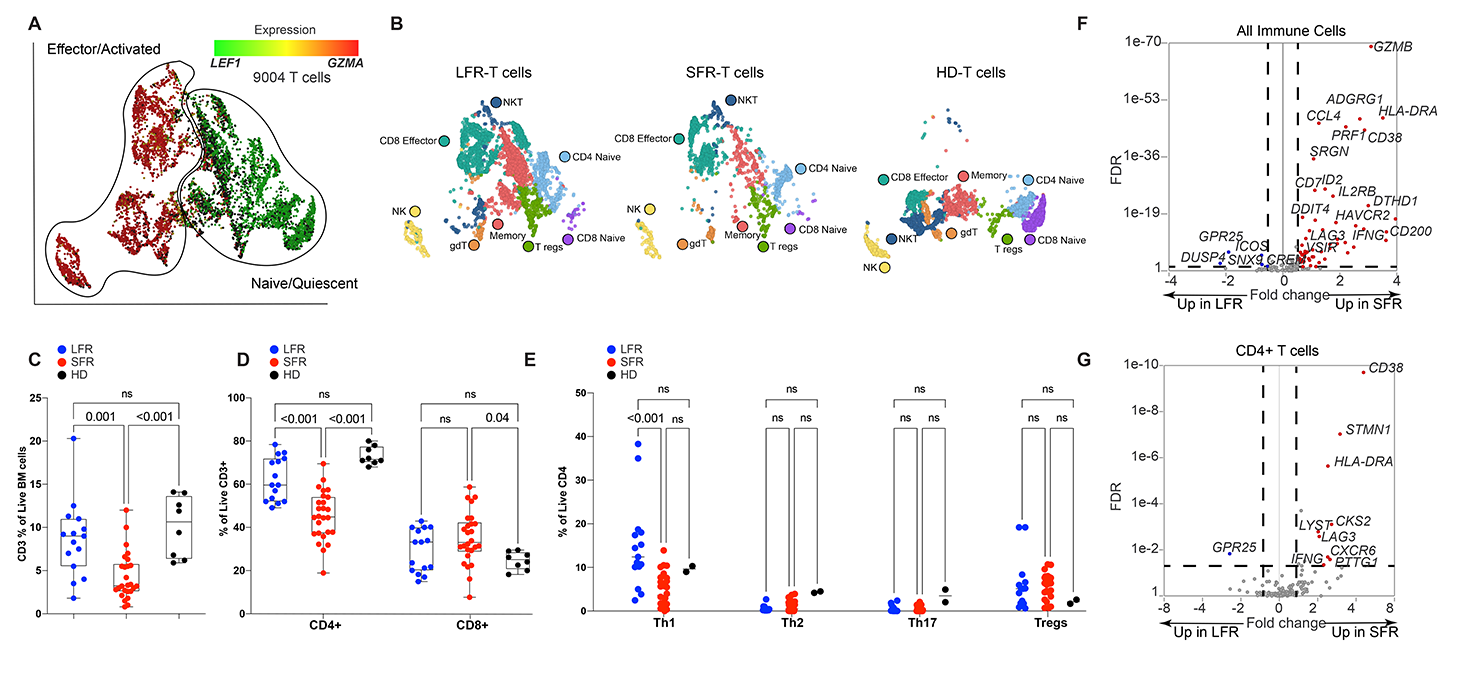
Single cell RNA-sequencing and flow cytometric studies of T cells from LFR vs. SFR cases. **Panels A and B** show UMAP projections of 9,004 T cells from the total bone marrow samples of two healthy donors and the 15 AML patients described in Figure 2. In **Panel A,** T cells are colored by the relative expression of *LEF1*, representing naïve/quiescent cells (green), vs. *GZMA*, representing activated/effector cells (red). In **Panel B,** T cell subsets are labeled by subtypes, identified by applying graph-based clustering, then identifying enriched biomarkers for each cluster. **Panels C through E:** plots displaying flow cytometric data from bone marrow samples from healthy donors (HD) or LFR vs. SFR cases, for **Panel C)** percentages of CD3+ cells, **Panel D)** percentages of CD4+ and CD8+ cells, and **Panel E)** percentages of Th1, Th2, Th17 and Treg subsets. Means were analyzed for significance with 2-way ANOVA and Tukey correction for multiple comparisons. **Panels F and G** display the results of an ANOVA comparison of 142 previously described Activation/Exhaustion markers^32,33^ in all 9,004 T cells (**Panel F**) and in CD4+ T cells (**Panel G**), comparing SFR to LFR cases. The X axis shows the log fold change for each gene, with no change (N/C) as the mid-point. The Y axis shows FDR in descending values. Differentially expressed genes (defined by log FC ± 1.3, FDR<0.05) are highlighted in red for SFR cases, or blue for LFRs.

To determine whether the proteins encoded by these immune checkpoint genes may be relevant for the suppression of T-cell function in AML marrows, we examined the expression of genes encoding known ligands for these receptors. MHC class II proteins serve as ligands for LAG3 (34), and as noted above, many MHC class II genes were more highly expressed in the SFR AML cells **(Figure 2D and Supplementary Figure 4D)**. *FGL1*, a recently described LAG3 ligand (35), was not expressed by AML cells (**Supplementary Figure 6A**). The gene encoding *PECAM1*, the ligand for CD38, was expressed similarly in both LFR and SFR AML cells (**Supplementary Figure 6B**). Of the known TIM3 ligands, the gene encoding *CEACAM* 1 was expressed in only a small fraction of AML cells (**Supplementary Figure 6C**), while *LGALS9* was more highly expressed in SFR AML cells (**Supplementary Figure 6E**, log FC = +1.3, FDR<0.001). Thus, when considering the potential relevance of immune checkpoint inhibition for relieving T-cell suppression in NK-AML, *LAG3* emerged as a leading candidate because of its high expression in SFR T-cells, and because the genes encoding its major ligands (MHC Class II) are highly expressed in the AML cells of most SFR cases.

### T-cell function in Long vs. Standard First Remission AML bone marrow samples

We next assessed the ability of AML cells from cryopreserved LFR and SFR bone marrow samples to influence the activation of their own infiltrating T-cells over 5 days *in vitro*, using CD3/CD28 stimulatory beads to activate the T-cell receptor (TCR). First, we evaluated unfractionated BM samples from LFR vs. SFR cases (i.e. containing both AML cells and T-cells) obtained at presentation. In the presence of AML cells, bead-activated CD4+ T-cells from the LFR cases significantly (p<0.001) upregulated the activation markers OX-40 and ICOS, while activation was blunted in the SFR cases (**Figure 4A and Supplementary Figure 7A**, graphs “*with AML”)*. LFR CD4 T-cells activated the LAG3 marker in the presence of AML cells (**Figure 4B**, p<0.001), while SFR cases had higher baseline percentages of CD4+LAG3+ cells (which corroborated the scRNA-seq data) that activated less robustly in the presence of AML cells (**Figure 4B**). PD-1 levels increased with T-cell stimulation, but there was no significant difference between LFR and SFR samples (**Supplementary Figure 7B**). We then tested the activation potential of CD3+ T-cells (purified by bead separation) from the same bone marrow samples; more than 95% of the AML cells were removed by this procedure. In the absence of AML cells, the activation of SFR CD4 T-cells was restored, and LAG3 baseline levels decreased to levels similar to that of LFR samples (**Figure 4A-B**, “*without AML”*). CD8+ cells displayed similar trends (**Supplementary Figure 7C-F**), although endogenous CD8+ T-cells suffered disproportionate losses in numbers after removal of AML cells, making measurements more difficult.

**Figure 4.**
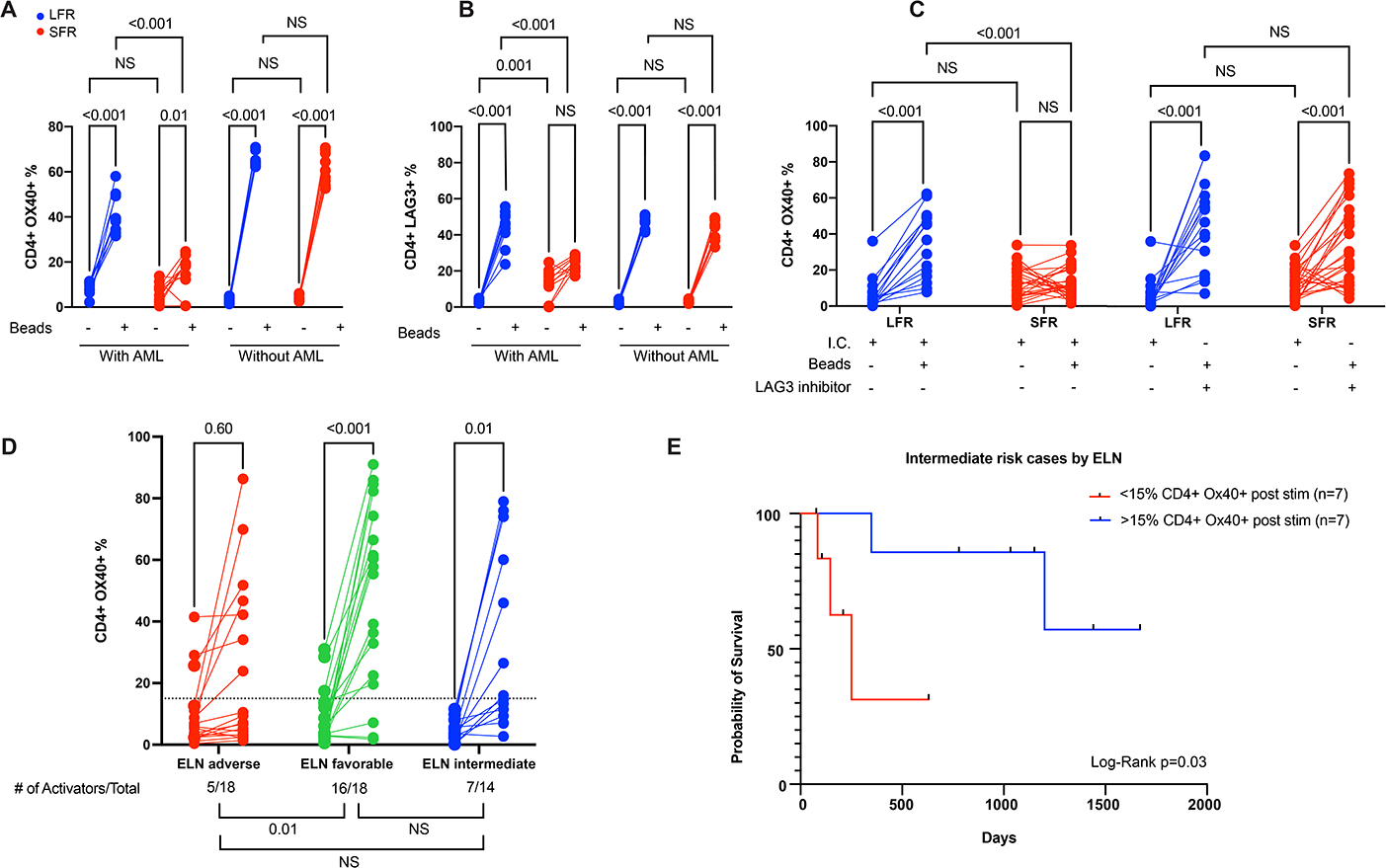
CD4+ T cell activation studies from presentation AML bone marrow samples. For the data shown in **Panels A through C,** cryovials from presentation bone marrow AML samples from LFR vs. SFR cases were thawed, and the fraction of CD3+ T cells was immediately defined by flow cytometry. These unfractionated samples were placed in media containing CD3/CD28 T cell receptor agonist beads in a 1:1 ratio with the previously defined number of T cells (“*with AML”*). An identical experiment was performed using bone marrow derived CD3+ T cells enriched from the same samples (“*without AML*”); CD3/CD28 beads were added at a 1:1 ratio after a 24 hour ‘washout’ period. T cell activation and inhibition markers were then quantified by flow cytometry 5 days later in both sets of experiments. The lines show the change in percentage of CD4+ cells expressing the activation marker OX-40 (**Panel A**), vs. the inhibitory marker LAG3 (**Panel B**) in samples treated with or without CD3/CD28 beads, which activate via the T cell receptor (as indicated by the legend below each graph). Red lines represent the SFR samples and blue lines represent LFR samples. 2-way ANOVA and Tukey multiple comparison tests were used to test for significance differences between groups. Results represent the summary of 3 independent experiments (n=10 unique samples from both the LFR and SFR sets). **Panel C** shows that similar levels of activation (as defined by OX40 expression) can be achieved in CD4+ T cells from SFR samples in the presence of a LAG3 blocking antibody at day 5 post stimulation. The negative controls (I.C.= Isotype Control) were treated with an isotype matched antibody. 2-way ANOVA and Tukey multiple comparison tests were used to test for significance differences between groups (n=15 for LFR and n=26 for SFR cases). **Panel D** shows changes in T cell activation, measured by the percentage of CD4+ cells expressing OX40, in 50 additional AML cases, representing an extension set. Unfractionated bone marrow samples from the presentation samples were evaluated 5 days after activation with CD3/CD28 T cell receptor agonist beads (in a 1:1 ratio with the number of measured T cells in each sample). The AML samples are grouped according to ELN category. Two-tailed t tests were used to calculate significance between pre- and post-activation samples. The threshold for CD4+ cell activation was defined as the median difference in activation (post stimulation over baseline) for the 50 samples tested with this assay (indicated by the black dotted line, at a fraction of15%). The number of AML samples that exhibited T cell activation in each ELN risk category, and a statistical comparison of the three groups, are shown below the graphs. Pearson’s Chi-squared test with Yates’ continuity correction was used to calculate differences in T cell activators among the groups. **Panel E** shows the relapse free survival for the intermediate ELN risk cases from the extension set (Panel D) stratified by CD4+ cell activation status. The blue and red lines represent samples with CD4+ cell activation above and below the activation threshold (defined as >15% CD4+ OX-40+ cells, and a fold change from baseline CD4+ OX-40 expression >=2.0). Vertical black lines indicate subjects at the time of censoring, as further defined in the Survival Analysis section of the Methods. Log rank (Mantel-Cox) test was used to estimate the difference in relapse free survival (p=0.03).

We repeated the 5-day CD3/CD28 bead stimulation protocol using unfractionated bone marrow samples from presentation (i.e. in the presence of AML cells), with or without a well-characterized LAG3 inhibitor (LAG3 blocking antibody, clone 17B4) (36, 37) or an isotype control antibody (“I.C.”). With LAG3 inhibition, endogenous CD4+ T-cell activation was restored in most SFR samples to levels that were similar to that of the LFR samples (**Figure 4C and Supplementary Figure 7G**).

To determine whether these findings were more broadly applicable, we performed the 5-day T-cell stimulation assay using presentation bone marrow samples from 50 additional *de novo* AML patients that were selected based on ELN 2017 classifications from DNA sequencing and cytogenetics, and on the availability of high quality cryovials from presentation bone marrow (see **Supplementary Table 11 and Supplementary Figure 8** for patient characteristics, mutational profiles, and ELN classifications). The fraction of activated (i.e. OX40+) CD4+ T-cells before and after stimulation for each sample are shown in **Figure 4D**. Activation was defined as >15% CD4+ OX-40+ cells, and a fold change from baseline CD4+ OX-40 expression >=2.0, which represented the median difference in activation (comparing post stimulation over baseline) for the 50 samples tested. 5/18 (28%) ELN adverse risk patients exhibited activation; RFS in these 5 cases was 1.2, 1.4, 4.5, 9.3, and 33.6 months. None of these patients wad transplanted in first remission. In contrast, 14/18 (78%) favorable risk patients exhibited CD4 activation. The strong correlation of CD4 activation with these ELN risk groups suggests that the mutational profile influences the immunosuppressive properties of AML cells--and can in fact supersede them in some cases. For intermediate risk cases, 7/14 (50%) displayed CD4 activation; importantly, the ‘activators’ in this risk group had significantly longer RFS (median survival not yet reached) compared with the non-activators (median survival of 250 days, Mantel-Cox Log-rank = 0.03, **Figure 4E**). These data suggest that T-cell activation induced by CD3/28 beads *in vitro* may provide novel prognostic information for the intermediate risk group that is not detected by current genetically-based classifications at presentation. Finally, we performed a multivariate analysis (using Firth logistic regression) to determine whether any covariate at presentation (including sex, age at diagnosis, white blood cell count, bone marrow blast percentage, ELN classification, or somatic mutations in AML-associated genes) was associated with CD4 activation status. Although ELN favorable risk patients were 8.18-times more likely to be CD4 activators, this difference did not reach significance due to wide confidence intervals. None of the other covariates were predictive of CD4 T-cell activation status at presentation (see **Supplementary Table 12**).

## Discussion

By evaluating a group of NK-AML patients with long first remissions after chemotherapy alone, we have been able to identify mechanisms that may be relevant for AML cell eradication after chemotherapy. Current prognostic tools, such as the ELN 2017 classification algorithm, have been especially helpful for guiding therapeutic decisions for favorable and adverse risk patients, but lose some resolution in the intermediate risk group, perhaps because these patients are more genetically heterogenous. The analysis of remission blood and bone marrow samples from LFR patients revealed that many do not appear to be living with persistent disease. Since a recent study showed that a small group of very late AML relapses (>5 years from diagnosis) arose from ancestral clones that were present at diagnosis (10), these observations suggest that eradication of virtually all AML cells after initial therapy is the basis for long term remissions for many AML patients.

Data from this study suggests that an immunosuppressive phenotype at presentation may negatively influence the effectiveness of standard chemotherapy. Specifically, we showed that bone marrow samples from LFR patients have well-preserved CD4 and CD8 populations that can react appropriately to TCR-mediated activation signals--even in the presence of their own AML cells. In contrast, SFR AML cells express high levels of MHC class II and genes associated with immune suppression. Their marrows have reduced numbers of immune cells (particularly of CD4+ Th1 cells) and their remaining T-cells often express an exhaustion signature (38). Importantly, T-cell activation was blocked in SFR samples by the presence of the AML cells themselves, and was reversible in most cases by removing the AML cells, or blocking the MHC class II inhibitory receptor, LAG3.

Several mechanisms could potentially explain these findings. For example, the AML cells of SFR cases may express mutations that encode neoantigens that cause prolonged antigen-specific T-cell activation, and eventual exhaustion (perhaps because these AML cells have developed mechanisms to resist T-cell mediated killing). However, using predictive algorithms, we did not detect substantial differences in putative neoantigen burdens or HLA alleles between LFR vs SFR cases (**Supplementary Figure 9A, B** and **Supplementary Table 13**). Alternatively, our data are consistent with a T-cell suppression phenotype in the SFR cases that may be mediated by the engagement of inhibitory T-cell receptors (i.e. checkpoints) by ligands that are expressed by AML cells; the specific mutations, genes, and pathways responsible for the expression of these inhibitory ligands will require further investigation. Additional mechanisms (e.g. inhibitory chemokines, metabolic inhibitors of T-cell function, and others) could also be relevant for this phenotype, and will need to be further explored.

Previous studies have shown that a dysfunctional CD4 T-cell population can dampen immune responses and lead to CD8 T-cell exhaustion (39–43); CD4 depleted tumors have decreased responsiveness to PD1 inhibition (44, 45) that may in part be dependent on MHC class II--LAG3 interactions, especially for tumors with high expression of MHC class II (like SFR AML cells) (46). Additionally, several groups have reported variable degrees of T-cell dysfunction in the bone marrow samples of AML patients (47–51), possibly caused by a suppressive effect of AML cells themselves (shown with AML cell lines (52), and also with primary AML samples (53)); here, we demonstrate that the absence of this immunosuppressive phenotype is strongly correlated with favorable clinical outcomes in both the LFR cases, and also in many ELN favorable and intermediate risk patients. Among patients with intermediate risk, RFS was significantly longer in samples with preserved T-cell activation. Importantly, the immune activation phenotype seems to be independent of other known prognostic factors in AML, such as age, ELN status, and/or somatic mutations. Remarkably, the concept that well preserved cell-mediated immunity can predict responses to chemotherapy in AML patients (both at diagnosis and during treatment) was proposed 50 years ago by Freireich and colleagues (13). In that study, AML patients with impaired delayed Type IV hypersensitivity skin testing, and/or impaired *in vitro* T-cell responses to mitogens, had higher rates of chemotherapy induction failure and relapse; interestingly, type IV reactions are now known to require the function of effector Th1 cells and macrophages (54), both of which appear to be altered in the SFR cases.

Although inhibition of PD1 and/or CTLA4 checkpoint inhibitors have not yet yielded clinical benefits for AML patients (55–57), this study suggests that alternative checkpoints may be relevant. We and others have identified LAG3, CD200 (24, 25, 58), and MRC1 (31) as potentially relevant targets for inhibition in AML. The means to inhibit these pathways currently exist, and are being explored in pre-clinical models and early phase clinical trials (59–61). The immunological phenotype of LFR patients strongly suggests that overcoming the immunosuppressive environment of ‘typical’ NK-AML cases at presentation could increase the chances for immunotherapy successes after conventional chemotherapy. The ability to integrate immunological phenotype assessment with current “up-front” risk stratification approaches—as Freireich et al. proposed with skin testing for Type IV hypersensitivity—may also prevent unnecessary allotransplantation, with its attendant complications. Future trials of immunologic modulation for the therapy of AML should therefore be guided both by the genotypes and phenotypes of AML cells—and of their immune microenvironments.

## Acknowledgements

Supported by a National Cancer Institute (NCI) K12 Program grant (CA167540) and a K08 grant (CA252632) to Dr. Ferraro, NCI Research Specialist Awards (R50 CA211466, to Dr. Rettig, and R50 CA211782, to Dr. Miller), the Genomics of Acute Myeloid Leukemia Program Project grant (P01 CA101937, to Dr. Ley), Edward P. Evans Foundation (to Dr. Walter), NCI Outstanding Investigator Awards (R35 CA197561, to Dr. Ley, and R35 CA210084 to Dr. DiPersio), a Barnes-Jewish Hospital Foundation grant (00335-0505-02, to Dr. Ley), a Specialized Program of Research Excellence in Acute Myeloid Leukemia grant (P50 CA171963, to Dr. Link), and NCI grants to the Alliance for banked samples from that entity (U10 CA180821, to Dr. Bertagnoli, U10 CA180882, to Dr. Mandrekar, and U24 CA196171, to Dr. Watson). We thank Dr. Allegra Petti for insightful discussions about the single-cell data analysis, and the Siteman Flow Cytometry Core, and the Tissue Procurement Core, supported by the Siteman Cancer Center support grant (P30 CA091842), and our patients for their participation. We dedicate this study to the memory of Dr. Clara Bloomfield.

