## Supplementary Appendix for "Immunosuppression and Outcomes in Acute Myeloid Leukemia"

### Supplementary Appendix Table of Contents

### A. Supplementary Results

#### A1. Long First Remission (LFR) Clinical Summaries (Washington University cases only)

#285. 61 year old female diagnosed with AML FAB subtype M1, following presentation with fatigue and bruising. Blood counts at presentation were WBC 25,500 cells/mm3, hemoglobin 7.9 g/dl, and platelets 57,000 cells/m3, with 96% circulating blasts. A bone marrow biopsy demonstrated 95% cellularity, with 91% blasts. Cytogenetics showed a 46 XX karyotype. Molecular diagnostic studies were not done. Initial therapy consisted of 7 + 3 induction with infusional cytarabinend idarubicin. A bone marrow biopsy at midcycle demonstrated an ablated marrow. A repeat marrow at count recovery demonstrated complete remission. Post-remission chemotherapy consisted of 2 cycles of intermediate dose cytarabine. She remains alive in remission, 17.8 years post-diagnosis.

#139406. 22 year old male diagnosed with AML (FAB indeterminate, with aberrant T cell antigen expression), following presentation with sore throat and cervical lymphadenopathy. Blood counts at presentation were WBC 36,300 cells/mm3, hemoglobin 12.1 g/dl, and platelets 95,000 cells/mm3, with 39% circulating blasts. A bone marrow biopsy demonstrated 80% cellularity, with 77% blasts. Cytogenetics showed a 46 XY karyotype. Molecular diagnostic study results included: FLT3 ITD positive, FLT3 D835 negative, NPM1 negative, and CEBPA negative. Initial therapy consisted of 7 + 3 induction with infusional cytarabine and daunorubicin. A bone marrow biopsy at midcycle demonstrated an ablated marrow. A repeat marrow at count recovery demonstrated complete remission. Post-remission chemotherapy consisted of 4 cycles of high dose cytarabine. He remains alive in remission, 9 years post-diagnosis.

#318748. 62 year old female diagnosed with AML FAB subtype M5a, following presentation with nausea, vomiting, and fatigue. Blood counts at presentation were WBC 94,300 cells/dl, hemoglobin 11.1 g/dl, and platelets 69,000 cells/dl, with 50% circulating blasts. A bone marrow biopsy demonstrated 90% cellularity, with 83% blasts. Cytogenetics showed a 46 XX karyotype. Molecular diagnostic study results included: FLT3 ITD positive, FLT3 D835 negative, and NPM1 positive. Initial therapy consisted of 7 + 3 induction with infusional cytarabine and idarubicin. Treatment was complicated by acute renal failure, sepsis, and bowel perforation. A bone marrow biopsy at midcycle demonstrated an ablated marrow. A repeat marrow at count recovery demonstrated complete remission. Post-remission chemotherapy consisted of 2 cycles of 5 + 2 infusional cytarabine and idarubicin. She remains alive in remission at last follow-up, 7.6 years post-diagnosis.

#334228. 63 year old male diagnosed with AML M1, following presentation with fatigue and anorexia. Blood counts at presentation were WBC 1,800 cells/mm3, hemoglobin 10.3g/dl, and platelets 13,000 cells/mm3, with 36% circulating blasts. A bone marrow biopsy demonstrated 80% cellularity, with 77% blasts. Cytogenetics showed a 46 XY karyotype. Molecular diagnostic study results included: FLT3 ITD negative, FLT3 D835 negative, NPM1 positive, and cKIT negative. Initial therapy consisted of 7 + 3 induction with cytarabine and idarubicin. A bone marrow biopsy at midcycle demonstrated an ablated marrow. A repeat marrow at count recovery demonstrated complete remission. Post-remission chemotherapy consisted of 3 cycles of intermediate dose cytarabine. He remains alive in remission at last follow-up, 5 years post-diagnosis.

#412761. 48 year old male diagnosed with AML FAB subtype M5, following presentation with leukocytosis and anemia. Blood counts at presentation were WBC 265,800 cells/mm3, hemoglobin

6.5 g/dl, and platelets 37,000 cells/mm3, with 40% circulating blasts, 13% “young monocytes,” and 30% monocytes. A bone marrow biopsy demonstrated >90% cellularity, with 83% blasts. Cytogenetics showed a 46 XY karyotype on peripheral blood. Molecular diagnostic study results included: FLT3 ITD negative and FLT3 D835 negative. Initial therapy consisted of emergent leukopheresis followed by 7 + 3 + 3 induction with infusional cytarabine, daunorubicin, and etoposide. A bone marrow biopsy at midcycle demonstrated an ablated marrow. A repeat marrow at count recovery demonstrated complete remission. Post-remission chemotherapy consisted of 3 cycles of high dose cytarabine. He remains alive in remission at last follow-up, 8 years post-diagnosis.

#452442. 47 year old male diagnosed with AML FAB subtype M2 following presentation with abscesses in his forearm and groin. Blood counts at presentation were WBC 57,700 cells/mm3, hemoglobin 7.5 g/dl, and platelets 10,000 cells/mm3, with 38% circulating blasts. A bone marrow biopsy demonstrated 90% cellularity, with 27% blasts. Cytogenetics showed a 46 XY karyotype. Molecular diagnostic study results included: FLT3 ITD negative, FLT3 D835 positive, and NPM1 positive. Past medical history was notable for rheumatoid arthritis treated with methotrexate. Initial therapy consisted of 7 + 3 induction with infusional cytarabine and idarubicin. A bone marrow biopsy at midcycle demonstrated an ablated marrow. A repeat marrow at count recovery demonstrated complete remission. Post-remission chemotherapy consisted of 3 cycles of intermediate dose cytarabine. He remains in complete remission at last follow-up, 7 years post-diagnosis.

#613389. 57 year old male diagnosed with AML FAB subtype M2, following presentation with bleeding after a hernia repair and incidentally noted anemia and thrombocytopenia. Blood counts at presentation were WBC 9,000 cells/mm3, hemoglobin 8.6 g/dl, and platelets 27,000 cells/mm3, with 49% circulating blasts. A bone marrow biopsy demonstrated 70% cellularity, with 25% blasts. Cytogenetics showed a 46 XY karyotype. Molecular diagnostic study results included: NPM1 negative, FLT3 ITD negative, and FLT3 D835 negative. Initial therapy consisted of 7 + 3 induction with infusional cytarabine and daunorubicin. A bone marrow biopsy at midcycle demonstrated an ablated marrow. A repeat marrow at count recovery demonstrated complete remission. Post-remission chemotherapy consisted of 4 cycles of high dose cytarabine. He remains alive in remission, 6.5 years post-diagnosis.

#627523. 19 year old male diagnosed with AML M4, following presentation with fatigue, anorexia, and sore throat. Blood counts at presentation were WBC 53,600 cells/mm3, hemoglobin 6.8 g/dl, and platelets 54,000 cells/mm3, with 6% circulating blasts and 50% monocytes. A bone marrow biopsy demonstrated 90% cellularity, with 74% blasts/promonocytes. Cytogenetics showed a 46 XY karyotype. Molecular diagnostic study results included: FLT3 ITD negative, FLT3 D835 positive, and NPM1 positive. Initial therapy consisted of 7 + 3 induction with infusional cytarabine and daunorubicin, with subsequent midostaurin (vs placebo). A bone marrow biopsy at midcycle demonstrated an ablated marrow. A repeat marrow at count recovery demonstrated complete remission. Post-remission chemotherapy consisted of 4 cycles of high dose cytarabine and subsequent midostaurin (vs placebo), followed by 12 months of maintenance midostaurin (vs placebo). He remains alive in remission, 8.7 years post-diagnosis.

#635258. 62 year old white male diagnosed with AML FAB subtype M5, following presentation with fever and dyspnea. Blood counts at presentation were WBC 101,800 cells/mm3, hemoglobin 9.5 g/dl, and platelets 38,000 cells/mm3, with 12% circulating blasts and 51% “young monocytes”. A bone marrow biopsy demonstrated 90% cellularity, with 85% blasts. Cytogenetics showed a 46 XY karyotype. Molecular diagnostic study results included: NPM1 positive, FLT3 ITD negative, and FLT3 D835 negative. Past medical history was notable for localized prostate cancer treated with radiation 2 years prior. Following cytoreduction with hydroxyurea, initial therapy consisted of 7 + 3 induction with infusional cytarabine and idarubicin. A bone marrow biopsy at midcycle demonstrated an ablated marrow. A repeat marrow at count recovery demonstrated complete remission. Post-remission chemotherapy consisted of 4 cycles of high dose cytarabine consolidation. He remains in complete remission, 12.5 years post-diagnosis.

#727185. 61 year old female diagnosed with AML FAB subtype M2, following presentation with recurrent diverticulitis, and incidentally noted leukocytosis and thrombocytopenia. Blood counts at presentation were WBC 16,400 cells/mm3, hemoglobin 12.0 g/dl, and platelets 55,000/mm3, with 10% circulating blasts. A bone marrow biopsy demonstrated 60-70% cellularity, with 23% blasts. Cytogenetics showed a 46 XX karyotype. Molecular diagnostic study results included: NPM1 negative, FLT3 ITD negative, and FLT3 D835 negative. Initial therapy consisted of 7 + 3 induction with infusional cytarabine and idarubicin. A bone marrow biopsy at midcycle demonstrated an ablated marrow. A repeat marrow at count recovery demonstrated complete remission. Post-remission chemotherapy consisted of 3 cycles of intermediate dose cytarabine. She remains alive in remission, 9.1 years post-diagnosis.

#759361. 54 year old female diagnosed with AML M2, following presentation with weakness and lightheadedness. Blood counts at presentation were WBC 3,300 cells/mm3, hemoglobin 8.7 g/dl, and platelets 18,000 cells/mm3, with “rare” circulating blasts. A bone marrow biopsy demonstrated 80% cellularity, with 42% blasts. Cytogenetics showed a 46 XX karyotype. Molecular diagnostic study results included: FLT3 ITD negative, FLT3 D835 negative, NPM1 positive, and BCR-ABL negative. Initial therapy consisted of 7 + 3 induction with infusional cytarabine and idarubicin. A bone marrow biopsy at midcycle demonstrated an ablated marrow. A repeat marrow at count recovery demonstrated complete remission. Post-remission chemotherapy consisted of 4 cycles of intermediate dose cytarabine. She remains alive in remission, 9.6 years post-diagnosis.

#807970. 38 year old male diagnosed with AML M1, following presentation with fatigue and cough. Blood counts at presentation were WBC 48,300 cells/mm3, hemoglobin 7.4 g/dl, and platelets 26,000 cells/mm3, with 99% circulating blasts. A bone marrow biopsy demonstrated >90% cellularity, with 86% blasts. Cytogenetics showed a 46 XY karyotype. Molecular diagnostic study results included: FLT3 ITD negative, FLT3 D835 negative, NPM1 positive, and BCR-ABL negative. Initial therapy consisted of 7 + 3 induction with infusional cytarabine and idarubicin. A bone marrow biopsy at midcycle demonstrated an ablated marrow. A repeat marrow at count recovery demonstrated complete remission. Post-remission chemotherapy consisted of 3 cycles of high dose cytarabine. He remains alive in remission, 12.9 years post-diagnosis.

#831711. 57 year old female diagnosed with AML FAB subtype M1, following presentation with fatigue and gum bleeding. Blood counts at presentation were WBC 4,400 cells/m3, hemoglobin 7.4 g/dl, and platelets 61,000 cells/mm3, with 38% circulating blasts. A bone marrow biopsy demonstrated 80% cellularity, with 64% blasts. Cytogenetics showed a 46 XX karyotype. Molecular diagnostic studies were not done. Following cytoreduction with hydroxyurea, initial therapy consisted of 7 + 3 induction with infusional cytarabine and idarubicin. A bone marrow biopsy at midcycle demonstrated an ablated marrow. A repeat marrow at count recovery demonstrated complete remission. Post-remission chemotherapy consisted of 3 cycles of high dose cytarabine. She remains in complete remission at last known follow-up, 14 years post-diagnosis. She was diagnosed with stage 1 invasive ductal breast carcinoma (ER+/PR+/HER2-), 14 years post-diagnosis of AML.

#854862. 41 year old female diagnosed with AML M2, following presentation with fatigue and shortness of breath. Blood counts at presentation were WBC 6,000 cells/mm3, hemoglobin 12.5 g/dl, and platelets 33,000 cells/mm3, with 20% circulating blasts. A bone marrow biopsy demonstrated 80% cellularity, with 42% blasts. Cytogenetics showed a 46 XX karyotype. Molecular diagnostic study results included: FLT3 ITD negative.  Initial therapy consisted of 7 + 3 induction with infusional cytarabine and idarubicin. A bone marrow biopsy at midcycle demonstrated an ablated marrow. A repeat marrow at count recovery demonstrated complete remission. Post-remission chemotherapy consisted of 4 cycles of high dose cytarabine. She remains alive in remission, 9.6 years post-diagnosis.

#868442. 52 year old male diagnosed with AML M4, following presentation with bilateral knee pain, HSV mucositis, and fever. Blood counts at presentation were WBC 329,200 cells/mm3, hemoglobin 7.9 g/dl, and platelets 70,000 cells/mm3, with 43% circulating blasts. A bone marrow biopsy demonstrated 90% cellularity, with 75% blasts. Cytogenetics showed a 46 XY karyotype. Molecular diagnostic study results included: FLT3 ITD negative, FLT3 D835 positive, NPM1 positive, and BCR-ABL negative. Following emergent leukopheresis and hydroxyurea, initial therapy consisted of 7 + 3 induction with infusional cytarabine and daunorubicin, with subsequent midostaurin (vs placebo). A bone marrow biopsy at midcycle demonstrated an ablated marrow. A repeat marrow at count recovery demonstrated complete remission. Post-remission chemotherapy consisted of 4 cycles of high dose cytarabine with subsequent midostaurin (vs placebo), followed by 12 months of maintenance midostaurin (vs placebo). He remains alive in remission, 10.6 years post-diagnosis.

#916462. 54 year old female diagnosed with AML FAB subtype M1, following presentation with fever and malaise. Blood counts at presentation were WBC 14,800 cells/mm3, hemoglobin 10.0 g/dl, and platelets 347,000 cells/mm3, with 65% circulating blasts. A bone marrow biopsy demonstrated 90% cellularity, with 83% blasts. Cytogenetics showed a 46 XY karyotype. Molecular diagnostic study results included: FLT3 ITD positive and JAK2 positive. Initial therapy consisted of 7 + 3 induction with infusional cytarabine and idarubicin. A bone marrow biopsy at midcycle demonstrated an ablated marrow. A repeat marrow at count recovery demonstrated complete remission. Post-remission chemotherapy consisted of 4 cycles of high dose cytarabine. She remains alive in complete remission, 9.6 years post-diagnosis.

#964973. 59 year old male diagnosed with AML M5b, following presentation with mouth sores. Blood counts at presentation were WBC 70,800 cells/mm3, hemoglobin 9.3 g/dl, and platelets 129,000 cells/mm3, with 5% circulating blasts and 59% monocytes. A bone marrow biopsy demonstrated 80% cellularity, with 83% blasts. Cytogenetics showed a 46 XX karyotype. Molecular diagnostic study results included: FLT3 ITD negative, FLT3 D835 negative, and NPM1 positive. Initial therapy consisted of 7 + 3 induction with infusional cytarabine and idarubicin. A bone marrow biopsy at midcycle demonstrated an ablated marrow. A repeat marrow at count recovery demonstrated complete remission. Post-remission chemotherapy consisted of 4 cycles of high dose cytarabine. He remains alive in remission at last known follow-up, 5.6 years post-diagnosis.

#973536. 56 year old male diagnosed with AML FAB subtype M4, following presentation with fatigue and dyspnea. Blood counts at presentation were WBC 26,600 cells/mm3, hemoglobin 9.1 g/dl, and platelets 31,000/mm3, with 7% circulating blasts and 37% monocytes. A bone marrow biopsy demonstrated 95% cellularity, with 34% blasts. Cytogenetics showed a 46 XY karyotype. Molecular diagnostic studies were not done. Initial therapy consisted of 7 + 3 induction with infusional cytarabine and idarubicin. A bone marrow biopsy at midcycle demonstrated an ablated marrow. A repeat marrow at count recovery demonstrated complete remission. Post-remission chemotherapy consisted of 4 cycles of high dose cytarabine. He remained in remission prior to succumbing to complications of non-alcoholic hepatic steatosis, 11 years post-diagnosis of AML.

#976838. 37 year old female diagnosed with AML FAB subtype M2, following presentation with weakness. Blood counts at presentation were WBC 2,400 cells/mm3, hemoglobin 10.6 g/dl, and platelets 43,000/mm3, without circulating blasts. A bone marrow biopsy demonstrated 80% cellularity, with 46% blasts. Cytogenetics showed a 46 XX karyotype. Molecular diagnostic study results included: FLT3 ITD negative, FLT3 D835 negative, and NPM1 positive. Initial therapy consisted of 4 + 3 induction with infusional cytarabine and idarubicin. A bone marrow biopsy at midcycle demonstrated an ablated marrow. A repeat marrow at count recovery demonstrated complete remission. Post-remission chemotherapy consisted of 4 cycles of 3 + 2 infusional cytarabine and idarubicin. She remains alive in remission, 5.7 years post-diagnosis of AML.

#### A2. Standard First Remission (SFR) Clinical Summaries

#287. 64 year old female diagnosed with AML FAB subtype M2, following presentation with fever. Blood counts at presentation were WBC 116,400 cells/mm3, hemoglobin 9.6 g/dl, and platelets 103,000/cells/mm3, with 5% circulating blasts and 42% promyelocytes. A bone marrow biopsy demonstrated 90% cellularity, with 23% blasts and 37% promyelocytes. Cytogenetics showed a 46 XX karyotype. FISH studies for PML/RAR rearrangement were negative. Molecular diagnostic studies were not done. Initial therapy consisted of 7 + 3 with infusional cytarabine and idarubicin. A bone marrow biopsy at midcycle demonstrated an ablated marrow. A repeat marrow following count recovery demonstrated complete remission. A lumbar puncture subsequently obtained to evaluate headache and altered mental status 52 days post-diagnosis demonstrated leukemic involvement, and she expired two weeks later without additional treatment.

#104851. 25 year old female diagnosed with AML FAB subtype M2, following presentation with excessive bleeding following wisdom teeth extraction. Blood counts at presentation were WBC 149,000 cells/mm3, hemoglobin 6.2 g/dl, and platelets 86,000 cells/mm3, with 84% circulating blasts. A bone marrow biopsy demonstrated 100% cellularity, with 53% blasts. Cytogenetics showed a 46 XX karyotype. Molecular diagnostic studies were not done. Following emergent leukopheresis and hydroxyurea, initial therapy consisted of 7 + 3 induction with infusional cytarabine and daunorubicin. A bone marrow biopsy at midcycle demonstrated an ablated marrow. A repeat marrow at count recovery demonstrated complete remission. Post-remission chemotherapy consisted of 3 cycles of high dose cytarabine. Relapse was documented 7 months post-diagnosis, and treated with mitoxantrone, etoposide, and intermediate dose cytarabine with disease persistence, and subsequent fludarabine, cytarabine, idarubicin, and gemtuzumab, following which a bone marrow biopsy showed ablation. She subsequently underwent 10/10 HLA-matched unrelated donor transplantation following myeloablative conditioning with cyclophosphamide, fludarabine, and total body irradiation, but expired due to post-transplant complications of pneumonia and sepsis at 11.5 months post-diagnosis.

#126620. 51 year old male diagnosed with AML M4, following presentation with fatigue and shortness of breath. Blood counts at presentation were WBC 124,000 cells/mm3, hemoglobin 3.9 g/dl, and platelets 41,000 cells/mm3, with 59% circulating blasts. A bone marrow biopsy demonstrated >95% cellularity, with 30-40% blasts/promonocytes. Cytogenetics showed a 46 XY karyotype. Molecular diagnostic study results included: NPM1 negative, FLT3 ITD positive, and FLT3 D835 negative. Initial therapy consisted of 7+3 induction with infusional cytarabine and idarubicin. A bone marrow biopsy at midcycle demonstrated an ablated marrow. A repeat marrow at count recovery demonstrated complete remission. Post-remission chemotherapy consisted of one cycle of intermediate dose cytarabine, following which he relapsed one month later. Salvage chemotherapy consisted of mitoxantrone, etoposide, and intermediate dose cytarabine, following which a repeat marrow biopsy showed persistent disease. He subsequently underwent 10/10 HLA-matched unrelated donor stem cell transplantation with active disease, following myeloablative conditioning with clofarabine and busulfan. Repeat marrow biopsies at approximately one, three, six, and twelve months post-transplant demonstrated complete remission with full donor engraftment. His post–transplant course was complicated by acute (skin) and extensive chronic (skin/mouth/eyes) graft versus host disease, and he remains alive 7.9 years post-transplant.

#161510. 44 year old male diagnosed with AML M2. Blood counts at presentation were WBC 12,800 cells/mm3, hemoglobin 8.4 g/dl, and platelets 74,000 cells/mm3, with 40% circulating blasts. A bone marrow biopsy demonstrated 90% cellularity, with 31% blasts. Cytogenetics showed a 46 XY karyotype. Molecular diagnostic study results included: JAK2 negative, BCR/ABL negative, NPM1 negative, FLT3 ITD negative, and FLT3 D835 negative. Initial therapy consisted of 7 + 3 induction with infusional cytarabine and idarubicin. A bone marrow biopsy at midcycle demonstrated 10-20% cellularity with 5-10% blasts, for which he was reinduced with 5 + 2. A repeat marrow at count recovery demonstrated persistent disease. He was subsequently treated with mitoxantrone, etoposide, and intermediate dose cytarabine without response. He subsequently underwent 10/10 HLA-matched unrelated donor stem cell transplantation with active disease, following myeloablative conditioning with busulfan and cyclophosphamide. He relapsed 2 months post-transplant, and was subsequently treated with decitabine without response; he expired 10 months post-diagnosis.

#186481. 57 year old female diagnosed with AML FAB subtype M4, following presentation with fatigue and skin nodules. Blood counts at presentation were WBC 110,200 cells/mm3, hemoglobin 12.0 g/dl, and platelets 72,000 cells/mm3, with “abundant” circulating blasts. A bone marrow biopsy demonstrated 80% cellularity, with 90% blasts. Cytogenetics showed a 46 XX karyotype. Molecular diagnostic studies were not done. A skin biopsy demonstrated leukemia cutis. Initial therapy consisted of 7 + 3 + 3 induction with infusional cytarabine, daunorubicin, and etoposide, and a concurrent investigational agent. A bone marrow biopsy at midcycle demonstrated an ablated marrow. A repeat marrow at count recovery demonstrated complete remission. Post-remission chemotherapy consisted of 2 cycles of intermediate dose cytarabine. Relapse was documented 5 months post-diagnosis, and despite salvage chemotherapy with mitoxantrone, etoposide, and intermediate dose cytarabine, she expired 2.5 months later.

#186706. 60 year old male diagnosed with AML FAB subtype M4, following presentation with weakness and fatigue. Blood counts at presentation were WBC 14,300 cells/mm3, hemoglobin 9.5 g/dl, and platelets 10,000 cells/mm3, with 20% circulating blasts, and 26% “young monocytes.” A bone marrow biopsy demonstrated 80% cellularity, with 77% blasts. Cytogenetics showed a 46 XY karyotype. Molecular diagnostic study results included: FLT3 ITD negative, D835 negative, and NPM1 negative. Initial therapy consisted of 7 + 3 induction with infusional cytarabine and idarubicin. A bone marrow biopsy at midcycle demonstrated persistent disease and he was re-induced with 5 + 2. Following a prolonged time to count recovery, a subsequent bone marrow biopsy demonstrated complete remission. Due to marginal performance status and co-morbidity, including COPD and coronary artery disease, he was deemed a poor candidate for intensive consolidation and was initially observed until a subsequent bone marrow biopsy demonstrated relapse at 4.5 months post-diagnosis. Salvage chemotherapy consisted of clofarabine and high dose cytarabine, following which a subsequent bone marrow biopsy demonstrated complete remission. Again, due to performance status and co-morbidity, he was not felt to be a candidate for allogeneic stem cell transplantation, and was enrolled in a study of oral clofarabine maintenance. Disease relapse was again documented 10 months post-diagnosis and treated with decitabine. He expired from complications of influenza infection prior to reassessment of disease status at 11 months post-diagnosis.

#208027. 62 year old female diagnosed with AML FAB subtype M2, initially managed with chlorambucil and hydroxyurea prior to emigration to the US and initial presentation at our center for treatment. Blood counts at presentation were WBC 75,200 cells/mm3, hemoglobin 11.1 g/dl, and platelets 110,000 cells/mm3, with 7% circulating blasts, and 11% "young monocytes." A bone marrow aspirate showed 46% blasts. Cytogenetics showed a 46 XX karyotype in 19/20 metaphases, with a single cell demonstrating an insertion at 7p22 of uncertain significance. Molecular diagnostic studies were not done. Initial therapy consisted of 7 + 3 induction with infusional cytarabine and idarubicin. A bone marrow biopsy at midcycle demonstrated an ablated marrow. A repeat marrow at count recovery demonstrated complete remission. Post-remission chemotherapy consisted of 3 cycles of high dose cytarabine. Relapse was documented 7 months post-diagnosis, which was observed initially and subsequently treated with high dose cytarabine and gemtuzumab ozogamycin. She expired 10 months post-diagnosis from complications of pneumonia, prior to re-evaluation of disease status.

#286032. 50 year old male diagnosed with AML M2, following presentation with weakness and dyspnea on exertion. Blood counts at presentation were WBC 18,900 cells/mm3, hemoglobin 6.1 g/dl, platelets 190,000 cells/mm3, with 20% circulating blasts. A bone marrow biopsy demonstrated 90% cellularity, with 32% blasts. Cytogenetics showed a 46 XY karyotype. Molecular diagnostic study results included: NPM1 positive, FLT3 ITD positive, and FLT3 D835 negative. Initial therapy consisted of induction with intermediate dose cytarabine and idarubicin. A bone marrow biopsy at midcycle demonstrated an ablated marrow. A repeat marrow at count recovery demonstrated complete remission. Post-remission chemotherapy consisted of 4 cycles of intermediate dose cytarabine and idarubicin. His disease relapsed 11 months post-diagnosis and was treated with mitoxantrone, etoposide, and intermediate dose cytarabine. A subsequent marrow showed complete remission, and he underwent 1 cycle of decitabine consolidation, prior to 9/10 HLA-mismatched unrelated donor stem cell transplantation, following myeloablative conditioning with busulfan and cyclophosphamide. His post-transplant course was complicated by GI GVHD. Isolated extramedullary relapse (thoracic spine and cerebrospinal fluid) was documented 3.9 years post-diagnosis, and treated with intrathecal chemotherapy and withdrawal of immunosuppression, complicated by a flare of GVHD. Relapse was again documented in marrow and cerebrospinal fluid at 4.7 years post-diagnosis and treated with gilteritinib and intrathecal chemotherapy. He again relapsed 5.4 years post-diagnosis and expired shortly thereafter on hospice.

#296361. 31 year old female diagnosed with AML M5a, following presentation with fatigue and sore throat. Blood counts at presentation were WBC 249,700 cells/mm3, hemoglobin 10.3 g/dl, platelets 64,000 cells/mm3, with 85% circulating monocytes. A bone marrow biopsy demonstrated >90% cellularity, with 83% monoblasts/promonocytes. Cytogenetics showed a 46 XX karyotype. Molecular diagnostic studies were not done. Initial therapy consisted of 7 +3 + 3 induction with infusional cytarabine, daunorubicin, and etoposide. A bone marrow biopsy at midcycle demonstrated an ablated marrow. A repeat marrow at count recovery demonstrated complete remission. Post-remission chemotherapy consisted of 1 cycle of intermediate dose cytarabine and etoposide, followed by autologous stem cell transplantation with busulfan and etoposide conditioning. Relapse was documented 6 months post-diagnosis and treated with fludarabine, intermediate dose cytarabine, idarubicin, and gemtuzumab ozagamycin. She expired prior to count recovery from complications of sepsis 8 months post-diagnosis.

#312340. 33 year old male diagnosed with AML FAB subtype M2, following presentation with nausea, vomiting, and chills. Blood counts at presentation were WBC 2800 cells/mm3, hemoglobin 7.3 g/dl, and platelets 61,000 cells/mm3, with no circulating blasts. A bone marrow biopsy was "hypocellular," with 33% blasts. Cytogenetics showed a 46 XY karyotype in 18 metaphases, with an isolated del 9q21 and del 20q12, respectively, in two additional clones, of uncertain significance. Molecular diagnostic studies were not done. Initial therapy consisted of 7 + 3 + 3 induction with infusional cytarabine, daunorubicin, and etoposide, and a concurrent investigational agent. A bone marrow biopsy at midcycle was equivocal, and a repeat marrow one week later showed ablation. A repeat marrow at count recovery demonstrated complete remission. Post-remission chemotherapy consisted of 1 cycle of intermediate dose cytarabine and etoposide, followed by autologous stem cell transplantation with busulfan and etoposide conditioning. Relapse was documented shortly after count recovery, at 5.7 months post-diagnosis, and treated with gemtuzumab ozogamycin prior to proceeding to 10/10 HLA-matched unrelated donor stem cell transplantation, following myeloablative conditioning with cyclophosphamide and total body irradiation. He expired one month post-transplant from complications of acute graft versus host disease, prior to reassessment of disease status.

#329614. 68 year old female diagnosed with AML FAB subtype M1, following presentation with chest pain and dyspnea on exertion. Blood counts at presentation were WBC 202,700 cells/mm3, hemoglobin 10.7 g/dl, and platelets 56,000 cells/mm3, with 88% circulating blasts. A bone marrow biopsy demonstrated 70% cellularity, with 90% blasts. Cytogenetics showed a 46 XX karyotype.

Molecular diagnostic study results included: FLT3 ITD positive and NPM1 positive. Following emergent leukopheresis and hydroxyurea, initial therapy consisted of 7 + 3 induction with infusional cytarabine and idarubicin. A bone marrow biopsy at midcycle demonstrated an ablated marrow. A repeat marrow at count recovery demonstrated complete remission. Post-remission chemotherapy consisted of 3 cycles of oral clofarabine. Relapse was documented 4 months post-diagnosis, and treated with decitabine for six cycles prior to relapse. She was treated subsequently with gemtuzumab ozogamycin and achieved remission, following which she was maintained on sorafenib until relapse 27 months post-diagnosis. She was briefly treated with carfilzomib without response, and subsequently with cladribine and cytarabine, again achieving remission, following which she was maintained on subcutaneous cytarabine and idarubicin until relapse. She was retreated with gemtuzumab ozogamycin and again maintained on decitabine until relapse, at which time she received no further treatment. She expired 3.1 years post-diagnosis.

#332131. 24 year old male diagnosed with AML FAB subtype M1, following presentation with weakness, fever, and epistaxis. Blood counts at presentation were WBC 223,800 cells/mm3, hemoglobin 9.6 g/dl, and platelets 103,000 cells/mm3. A manual differential demonstrated 76% circulating blasts. A bone marrow biopsy demonstrated >90% cellularity, with 77% blasts. Cytogenetics showed a 46 XY karyotype. Molecular diagnostic study results included: FLT3 ITD positive and NPM1 negative. Following emergent leukopheresis and hydroxyurea, initial therapy consisted of 7 + 3 + 3 induction with infusional cytarabine, daunorubicin, and etoposide. A bone marrow biopsy at midcycle demonstrated an ablated marrow. A repeat marrow following count recovery demonstrated complete remission. Post-remission chemotherapy consisted of 3 cycles of high dose cytarabine. Relapse was documented in cerebrospinal fluid and bone marrow 10 months post-diagnosis, and treated with mitoxantrone, etoposide, and intermediate dose cytarabine. He subsequently underwent 6/6 HLA-matched sibling donor allogeneic stem cell transplantation, following myeloablative conditioning with cyclophosphamide and total body irradiation. He remained in remission, but expired from transplant complications of thrombotic thrombocytopenia purpura and sepsis 20.5 months post-diagnosis.

#375182. 57 year old male diagnosed with AML FAB subtype M5, following incidental discovery of leukocytosis during workup for back surgery. Blood counts at presentation were WBC 99,200 cells/mm3, hemoglobin 9.8 g/dl, and platelets 80,000 cells/mm3, with 4% circulating blasts. A bone marrow biopsy demonstrated >90% cellularity, with 52% blasts. Cytogenetics showed a 46 XY karyotype. Molecular diagnostic study results included: FLT3 D835 positive and FLT3 ITD negative. Initial therapy consisted of 7 + 3 induction with infusional cytarabine and idarubicin. A bone marrow biopsy at midcycle demonstrated an ablated marrow. A repeat marrow at count recovery demonstrated complete remission. Post-remission chemotherapy consisted of 3 cycles of high dose cytarabine. Relapse was documented 9 months post-diagnosis, and he expired from intracranial hemorrhage prior to receiving further therapy.

#433325. 51 year old female diagnosed with AML FAB subtype M2, following presentation with a urinary tract infection. Blood counts at presentation were WBC 32,900 cells/mm3, hemoglobin 13.5 g/dl, and platelets 194,000 cells/mm3, with 61% circulating monocytes. A bone marrow biopsy demonstrated 90% cellularity, with 64% blasts. Cytogenetics showed a 46 XX karyotype. Molecular diagnostic studies were not done. Initial therapy consisted of 7 + 3 + 3 induction with infusional cytarabine, daunorubicin, and etoposide. A bone marrow biopsy at midcycle demonstrated an ablated marrow. A repeat marrow at count recovery demonstrated complete remission. Post-remission chemotherapy consisted of 3 cycles of high dose cytarabine. Relapse was documented 10 months post-diagnosis, and treated with fludarabine, intermediate dose cytarabine, and idarubicin. She subsequently underwent 10/10 HLA-matched unrelated donor transplantation with active disease following myeloablative conditioning with cyclophosphamide and total body irradiation. Relapse was again documented 2 months post-transplant, and she expired 16 months post-diagnosis.

#508084. 38 year old male diagnosed with AML M4 following presentation with cellulitis and dyspnea. Blood counts at presentation were WBC 140,100 cells/mm3, hemoglobin 7.3 g/dl, and platelets 53,000 cells/mm3, with 53% circulating blasts. A bone marrow biopsy demonstrated 80-90% cellularity, with 53% blasts/promonocytes. Cytogenetics showed a 46 XY karyotype. Molecular diagnostic study results included: FLT3 ITD positive, FLT3 D835 negative, NPM1 negative, IDH1 negative, IDH2 negative, and DNMT3A (R882) negative. Initial therapy consisted of 7 + 3 induction with infusional cytarabine and idarubicin. A bone marrow biopsy at midcycle demonstrated an ablated marrow. A repeat marrow at count recovery demonstrated persistent disease. Salvage chemotherapy consisted of mitoxantrone, intermediate dose cytarabine, and etoposide. A bone marrow biopsy at count recovery again showed persistent disease, and he was treated with fludarabine, intermediate dose cytarabine, and idarubicin, followed three weeks later by 3/6 HLA-matched haploidentical stem cell transplantation with myeloablative fludarabine and fractionated total body irradiation conditioning, and post-transplant cyclophosphamide. He remains alive in remission 5.2 years post-diagnosis.

#509754. 21 year old female, diagnosed with AML M1, following presentation with fever and upper respiratory symptoms. Blood counts at presentation were WBC 8200 cells/mm3, hemoglobin 7.9 g/dl, and platelets 52,000 cells/mm3, with 53% circulating blasts. A bone marrow biopsy demonstrated 95% cellularity, with 91% blasts. Cytogenetics showed a 46 XX karyotype. FISH studies for a PML/RAR fusion were negative. Molecular diagnostic study results were not done. Initial therapy consisted of 7 + 3 + 3 induction with infusional cytarabine, daunorubicin, and etoposide. A bone marrow biopsy at midcycle demonstrated 9% blasts with <10% cellularity. A repeat marrow at count recovery demonstrated complete remission, however relapse was documented 3 weeks later, prior to consolidation therapy. She was treated with a salvage regimen of mitoxantrone, etoposide, and intermediate dose cytarabine, following which a repeat marrow demonstrated complete remission.

She subsequently underwent allogeneic 6/6 HLA-matched sibling donor stem cell transplantation, following myeloablative conditioning with cyclophosphamide and total body irradiation. She remains in alive and in remission 15.7 years post-diagnosis.

#606061. 37 year old female diagnosed with AML FAB subtype M2, following presentation with fatigue and easy bruisability. Blood counts at presentation were WBC 22,900 cells/mm3, hemoglobin 11.2 g/dl, and platelets 55,000 cells/mm3. A manual differential demonstrated 41% circulating blasts. A bone marrow biopsy demonstrated 90% cellularity, with 52% blasts. Cytogenetics showed a 46 XX karyotype. Molecular diagnostic studies were not done. Initial therapy consisted of 7 + 3 + 3 induction with infusional cytarabine, daunorubicin, and etoposide. She underwent a lumbar puncture at diagnosis for evaluation of headache and received a single dose of intrathecal methotrexate. CSF studies were negative for malignancy. A bone marrow biopsy at midcycle demonstrated an ablated marrow. A repeat marrow at count recovery demonstrated remission. Post-remission chemotherapy consisted of 1 cycle of intermediate dose cytarabine and etoposide, followed by autologous stem cell transplant with busulfan and etoposide conditioning. Relapse was documented 8.5 months post-diagnosis, and treated with mitoxantrone, etoposide, and cytarabine. She developed mucor mycosis infection shortly thereafter and expired 9 months post-diagnosis.

#721214. 41 year old female diagnosed with AML M1, following presentation with leukocytosis. Blood counts at presentation were WBC 151,800 cells/mm3, hemoglobin 12.3 g/dl, and platelets 46,000 cells/mm3, with 19% circulating blasts. A bone marrow biopsy demonstrated 90% cellularity, with 92% blasts. Cytogenetics were non-informative due to lack of dividing cells. Molecular diagnostic study results included: FLT3 ITD positive, FLT3 D835 negative, and NPM1 positive. Initial therapy consisted of 7 + 3 induction with infusional cytarabine and idarubicin. A bone marrow biopsy at midcycle demonstrated persistent disease. Salvage chemotherapy consisted of fludarabine, intermediate dose cytarabine, and idarubicin. A subsequent bone marrow biopsy showed persistent disease, and she treated with mitoxantrone, intermediate dose cytarabine, and etoposide,with a concurrent investigational agent, following which she had rapid disease progression at count recovery. She expired with persistent leukemia 5 months post-diagnosis.

#723101. 66 year old male diagnosed with AML FAB subtype M1, following presentation with fatigue and low grade fever. Blood counts at presentation were WBC 143,000 cells/mm3, hemoglobin 9.4 g/dl, and platelets 247,000 cells/mm3, with 73% circulating blasts. A bone marrow biopsy demonstrated 90% cellularity, with 79% blasts. Cytogenetics showed a 46 XY karyotype. Molecular diagnostic study results included: NPM1 negative and FLT3 ITD positive. Following emergent leukopheresis, initial therapy consisted of 7 + 3 induction with infusional cytarabine and idarubicin. A bone marrow biopsy at midcycle demonstrated persistent disease. Salvage chemotherapy consisted of mitoxantrone, etoposide, and intermediate dose cytarabine. A repeat marrow at count recovery demonstrated complete remission, however a bone marrow biopsy one month later showed disease relapse. Subsequent salvage chemotherapy consisted of cladribine, intermediate dose cytarabine, mitoxantrone, and filgrastim, with persistent disease, and he proceeded to 10/10 HLA-matched unrelated donor stem cell transplantation with active disease, following myeloablative conditioning with cyclophosphamide and total body irradiation. A bone marrow at 1 month post-transplant demonstrated remission, but he relapsed at 2 months post-transplant, he expired 6 months post-diagnosis.

#823477. 51 year old male diagnosed with AML M0, following presentation with diffuse musculoskeletal pain and blurred vision. Blood counts at presentation were WBC 10,900 cells/mm3, hemoglobin 9.8 g/dl, and platelets 129,000 cells/mm3, with 44% circulating blasts. A bone marrow biopsy demonstrated >90% cellularity, with “sheets of immature mononuclear precursors”. Cytogenetics showed a 46 XY karyotype. Molecular diagnostic studies were notable for: NPM1 negative, FLT3 ITD positive, and FLT3 D835 negative. Initial therapy consisted of 7 + 3 induction with infusional cytarabine and idarubicin. A bone marrow biopsy at midcycle demonstrated an ablated marrow. A repeat marrow at count recovery demonstrated complete remission. Post-remission chemotherapy consisted of 4 cycles of idarubicin and intermediate dose cytarabine. Relapse was documented in both bone marrow and cerebrospinal fluid 8 months post-diagnosis and treated with mitoxantrone, etoposide, and intermediate dose cytarabine salvage with concurrent intrathecal chemotherapy. He subsequently underwent 6/6 HLA-matched sibling donor allogeneic stem cell transplant, following myeloablative conditioning with cyclophosphamide and total body irradiation in second complete remission. Second relapse was documented 16 months post-diagnosis and treated with fludarabine, intermediate dose cytarabine, and idarubicin salvage chemotherapy, prior to second allogeneic transplant with a 3/6 HLA-matched haploidentical donor following myeloablative fludarabine, busulfan, and cyclophosphamide conditioning, followed by post-transplant cyclophosphamide. Isolated central nervous system relapse was documented in cerebrospinal fluid 23 months post-diagnosis and treated with craniospinal radiation and subsequent intrathecal chemotherapy. He subsequently received 4 cycles of decitabine maintenance prior to isolated skin relapse 32 months post-diagnosis, treated with gemtuzumab ozagamycin. He expired from progressive disease 3 years post-diagnosis.

#869586. 23 year old diagnosed with AML FAB subtype M4, following presentation with a ruptured appendix. Blood counts at presentation were WBC 27,100 cells/mm3, hemoglobin 12.0 g/dl, and platelets 23,000 cells/mm3, with 63% circulating blasts. A bone marrow biopsy was “inevaluable” for cellularity, with 51% blasts. Cytogenetics showed a 46 XY karyotype. Molecular diagnostic studies were not done. Initial therapy consisted of 7 + 3 + 3 induction with infusional cytarabine, daunorubicin, and etoposide. A bone marrow biopsy at midcycle demonstrated an ablated marrow. A repeat marrow at count recovery demonstrated complete remission. Post-remission therapy consisted of intermediate dose cytarabine and etoposide, followed by autologous stem cell transplantation following conditioning with busulfan and cyclophosphamide. Relapse was documented seven months post-diagnosis, and treated with fludarabine, intermediate dose cytarabine, idarubicin, and gemtuzumab ozogamycin with concurrent filgrastim, resulting in second complete remission. He subsequently underwent 6/6 HLA-matched sibling donor allogeneic stem cell transplantation, following myeloablative conditioning with cyclophosphamide and total body irradiation, but relapsed two months later. Subsequent treatment included: clofarabine and intermediate dose cytarabine; decitabine; and cladribine, intermediate dose cytarabine, and imatinib, as well as serial donor lymphocyte infusions, without achieving remission. He expired from progressive disease 19 months post-diagnosis.

#869922. 56 year old female diagnosed with AML FAB subtype M2, following presentation with fever and rash. Blood counts at presentation were WBC 202,000 cells/mm3, hemoglobin 9.9 g/dl, and platelets 45,000 cells/mm3, with 96% circulating blasts. A bone marrow biopsy demonstrated 80% cellularity, with 60% blasts. Cytogenetics showed a 46 XX karyotype. Molecular diagnostic studies were not done. Following emergent leukopheresis and hydroxyurea, initial therapy consisted of 7 + 3 + 3 induction with infusional cytarabine, daunorubicin, and etoposide. A bone marrow biopsy at midcycle demonstrated an ablated marrow. A repeat marrow at count recovery demonstrated complete remission. Post-remission chemotherapy consisted of 1 cycle of intermediate dose cytarabine and etoposide, followed by autologous stem cell transplant following conditioning with busulfan and cyclophosphamide, and post-transplant decitabine maintenance. Relapse was documented 9 months post-diagnosis, and treated with mitoxantrone, etoposide, and intermediate cytarabine. She subsequently underwent 10/10 HLA-matched unrelated donor transplantation following reduced intensity conditioning with fludarabine, busulfan, and thymoglobulin in second complete remission, but relapsed 2 months later, and expired on hospice 16 months post-diagnosis.

#875663. 68 year old female diagnosed with AML M4, following presentation with weakness and shortness of breath. Blood counts at presentation were WBC 57,700 cells/mm3, hemoglobin 10.0

g/dl, platelets 59,000 cells/mm3, with 25% circulating monocytes. A bone marrow biopsy demonstrated 80% cellularity, with 59% blasts. Cytogenetics showed a 46 XX karyotype (5 metaphases). Molecular diagnostic study results included: NPM1 positive, FLT3 ITD negative, and FLT3 D835 positive. Initial therapy consisted of 7 + 3 with infusional cytarabine and daunorubicin, with concurrent sorafenib. A bone marrow biopsy at midcycle demonstrated an ablated marrow. A repeat marrow at count recovery demonstrated complete remission. Post-remission chemotherapy consisted of 2 cycles of intermediate dose cytarabine with concurrent sorafenib and subsequent sorafenib maintenance, until relapse 12 months post-diagnosis. She was subsequently treated with an investigational anti-CD33 antibody without response, followed by decitabine without response, and azacytidine without response. She then underwent 9/10 -HLA-mismatched unrelated donor stem cell transplantation following reduced intensity conditioning with fludarabine, busulfan, and thymoglobulin. A bone marrow biopsy one-month post-transplant demonstrated complete remission with 92-97% donor engraftment. A repeat marrow two months later demonstrated relapse, for which she was treated with mitoxantrone, intermediate dose cytarabine, and etoposide, and a subsequent donor lymphocyte infusion. A midcycle bone marrow biopsy demonstrated an ablated marrow, however three weeks later circulating blasts were observed in peripheral blood, for which no further therapy was given, and she expired shortly thereafter, at 22 months post-diagnosis.

#992966. 67 year old male diagnosed with AML FAB subtype M4, following presentation with bone pain. Blood counts at presentation were WBC 47,400 cells/mm3, hemoglobin 8.3 g/dl, and platelets 154,000 cells/mm3, with 45% circulating blasts and 19% “young monocytes.” A bone marrow biopsy demonstrated 90% cellularity, with 89% blasts. Cytogenetics showed a 46 XY karyotype. Molecular diagnostic study results included: FLT3 negative. Initial therapy consisted of 7 + 3 induction with infusional cytarabine and daunorubicin. A bone marrow biopsy at midcycle demonstrated persistent disease, and he was re-induced with 5 + 2. A repeat marrow at count recovery demonstrated complete remission. Post-remission chemotherapy consisted of 1 cycle of intermediate dose cytarabine. Relapsed disease was documented 4.5 months post-initial diagnosis, and he expired 1 month later.

#115225. 22 year old male diagnosed with AML (FAB indeterminate), following presentation with fatigue, dyspnea on exertion, and fever. Blood counts at presentation were WBC 174,800 cells/mm3, hemoglobin 6.4 g/dl, and platelets 82,000 cells/mm3, with 94% circulating blasts. A bone marrow biopsy demonstrated >90% cellularity, with “sheets of blasts”. Cytogenetics showed a 46 XY karyotype. Molecular diagnostic study results included: NPM1 negative, FLT3 ITD positive, and FLT3 D835 negative. Initial therapy consisted of 7 + 3 induction with infusional cytarabine and daunoubicin, with subsequent midostaurin (vs. placebo). A bone marrow biopsy at midcycle demonstrated an ablated marrow. A repeat marrow at count recovery demonstrated complete remission. Post-remission chemotherapy consisted of 3 cycles of high dose cytarabine with subsequent midostaurin (vs placebo), followed by 12 months of maintenance midostaurin (vs placebo). Relapse was documented 28 months post-diagnosis and treated with mitoxantrone, etoposide, and intermediate dose cytarabine, with concurrent plerixafor, resulting in second complete remission. He subsequently underwent 3/6 matched (haploidentical) related donor allogeneic stem cell transplantation following myeloablative conditioning with fludarabine, cyclophosphamide, and total body irradiation, with post-transplant cyclophosphamide. He remains alive in remission approximately 10 years post-diagnosis.

#141273. 64 year old female diagnosed with AML M2, following presentation with fatigue and easy bruisability. Blood counts at presentation were WBC 70,300 cells/mm3, hemoglobin 10.6 g/dl, and platelets 35,000 cells/mm3, with 82% circulating blasts. A bone marrow biopsy demonstrated 95% cellularity, with 80% blasts. Cytogenetics showed a 46 XX karyotype. Molecular diagnostic study results included: NPM1 positive, FLT3 ITD positive, and FLT3 D835 negative. Initial therapy consisted of 7 + 3 induction with infusional cytarabine and idarubicin. A bone marrow biopsy at midcycle demonstrated an ablated marrow. A repeat marrow at count recovery demonstrated complete remission. Post-remission chemotherapy consisted of 3 cycles of high dose cytarabine with concurrent ATRA. Relapse was documented 15 months post-diagnosis and treated with cladribine, intermediate dose cytarabine, midostaurin, and ATRA with achievement of remission. Her disease relapsed 8 months later was treated with mitoxantrone, etoposide, and intermediate dose cytarabine. She expired from relapsed disease 2.3 years post-diagnosis.

#150288. 51 year old male diagnosed with AML M1, following presentation with fever and upper respiratory congestion. Blood counts at presentation were WBC 1,900 cells/mm3, hemoglobin 11.1 g/dl, and platelets 44,000 cells/mm3, without circulating blasts. A bone marrow biopsy demonstrated 30-60% cellularity with 62% blasts. Cytogenetics showed a 46 XY karyotype. Molecular diagnostic study results included: NPM1 negative, FLT3 ITD negative, and FLT3 D835 negative. Initial therapy consisted of 7 + 3 induction with infusional cytarabine and daunorubicin. A bone marrow biopsy at midcycle demonstrated an ablated marrow. A repeat marrow at count recovery demonstrated complete remission. Post-remission chemotherapy consisted of 3 cycles of high dose cytarabine. Relapse was documented 14 months post-diagnosis and treated with mitoxantrone, etoposide, and intermediate dose cytarabine, with concurrent plerixafor, resulting in second complete remission. He subsequently underwent 6/6 HLA-matched sibling donor allogeneic stem cell transplantation, following myeloablative conditioning with busulfan and cyclophosphamide. His disease relapsed again 18 months post-diagnosis and was treated with cladribine, intermediate dose cytarabine, midostaurin, and ATRA without response, and he expired 23 months post-diagnosis.

#311636. 64year old female diagnosed with AML M4, following presentation with fatigue, rectal bleeding, and easy bruisability. Blood counts at presentation were WBC 131,500 cells/m3, hemoglobin 10.2 g/dl, and platelets 15,000 cells/mm3, with 90% monocytes. A bone marrow biopsy demonstrated 90% cellularity, with 72% blasts. Cytogenetics showed a 46 XX karyotype. Molecular diagnostic studies were not done. Initial therapy consisted of 7 + 3 induction with infusional cytarabine and mitoxantrone, complicated by intracranial hemorrhage which was managed conservatively. A bone marrow biopsy at midcycle demonstrated an ablated marrow. A repeat marrow at count recovery demonstrated complete remission. Post-remission chemotherapy consisted of 2 cycles of intermediate dose cytarabine. Relapse was documented 12 months post-diagnosis and treated with mitoxantrone, etoposide, and intermediate dose cytarabine with no response, and subsequently with gemtuzumab ozogamycin.  She expired on hospice 2 years post-diagnosis.

#418499. 35 year old male diagnosed with AML M4, following presentation with increasing dyspnea on exertion. Blood counts at presentation were WBC 52,900 cells/mm3, hemoglobin 8.3 g/dl, and platelets 44,000 cells/mm3, with 18% blasts and 33% monocytes. A bone marrow biopsy demonstrated 100% cellularity, comprised of “almost entirely Leder-negative blasts”. Cytogenetics showed a 46 XY karyotype. Molecular diagnostic studies were not done. Initial therapy consisted of 7 + 3 induction with infusional cytarabine and idarubicin. A bone marrow biopsy at midcycle demonstrated persistent disease, and he was re-induced with high dose cytarabine. A repeat marrow at count recovery demonstrated complete remission. Post-remission chemotherapy consisted of 1 cycle of high dose cytarabine. He subsequently presented with a dental abscess and relapsed disease, and expired prior to additional therapy from complications of myocardial infarction, 17 months post-diagnosis.

#452198. 55 year old male diagnosed with AML M5, following presentation with fatigue and weight loss. Blood counts at presentation were WBC 72,600 cells/mm3, hemoglobin 8.2 g/dl, and platelets 17,000 cells/mm3, with 8% circulating blasts and 50% monocytes. A bone marrow biopsy was inevaluable for cellularity. The aspirate demonstrated 97% monoblasts. Cytogenetics showed a 46 XY karyotype. Molecular diagnostic studies were not done. Following cytoreduction with hydroxyurea, initial therapy consisted of 7 + 3 induction with infusional cytarabine and idarubicin. A bone marrow biopsy at midcycle demonstrated an ablated marrow. A repeat marrow at count recovery demonstrated complete remission. Post-remission therapy consisted of 4 cycles of intermediate dose cytarabine. Relapse was documented 16 months post-diagnosis, and treated with mitoxantrone, etoposide, and intermediate dose cytarabine with concurrent plerixafor. He underwent 6/6 HLA-matched sibling donor stem cell transplantation in second complete remission following myeloablative conditioning with busulfan and cyclophosphamide, but relapsed 108 months post-diagnosis. He was re-induced with 7 + 3 chemotherapy but expired due to sepsis prior to re-evaluation for response 9 years post-diagnosis.

#906708. 75 year old female diagnosed with AML M4, following presentation with fatigue and dyspnea. Blood counts at presentation were WBC 5,000 cells/mm3, hemoglobin 10.3 g/dl, and platelets 52,000 cells/mm3, with 16% circulating blasts. A bone marrow biopsy demonstrated 90% cellularity, with 91% blasts. Cytogenetics showed a 46 XX karyotype. Molecular diagnostic study results included: FLT3 ITD negative and FLT3 D835 negative. Initial therapy consisted of 7 + 3 induction with infusional cytarabine and idarubicin. A bone marrow biopsy at midcycle demonstrated an ablated marrow. A repeat marrow biopsy following prolonged time to count recovery showed no evidence of disease, but demonstrated complex cytogenetics, for which she was treated with 5+2 reinduction. Subsequent biopsies showed a persistent complex karyotype but no evidence of disease, until relapse with AML and background MDS was documented 17 months post-diagnosis, and treated with arsenic and decitabine with no response. She expired 2 years post-diagnosis.

#### B. Patient selection criteria

##### B1. Long First Remission selection

Inclusion Criteria:

1) morphologically documented *de novo* AML, with adequate bone marrow and skin DNA, and bone marrow RNA, for sequencing studies, 2) at least 18 years of age, 3) normal cytogenetics at presentation, 4) received “7+3” (or similar variants) for induction chemotherapy, 5) received high dose AraC as the primary consolidation therapy, and 6) the first remission lasted at least 5 years.

Of the **1,579** banked AML samples available at the time of sample selection, only **19 (1.2%)** met the inclusion criteria and had adequate material for analysis. All remain in continuous remission at this writing, with follow up times of 5.4-17 years.

Eight of these 19 cases had samples banked in remission, which allowed for assessment of mutation persistence in remission (see **Table 2**). Eleven of these cases had material banked only at presentation (because serial banking of AML cases started in 2011 in our study). Identical inclusion parameters were used to identify **9 additional cases from CALGB/Alliance studies from a total of 846 NK-AML** samples that met inclusion criteria **(1.1%).** Only bone marrow DNAs from the presentation samples were available for these cases. Informed consent for sample and data sharing, and genetic studies was obtained for all patients.

#### B2. Standard First Remission selection

Inclusion Criteria:

1) morphologically documented *de novo* AML, with adequate bone marrow and skin DNA, and bone marrow RNA, for sequencing studies, 2) at least 18 years of age, 3) normal cytogenetics at presentation, 4) received 7+3 induction and high doses AraC consolidation (or similar variation chemotherapies) 5) primary refractory (defined as failure to achieve remission after induction) or documented morphologic relapse within 24 months of achieving remission.

#### B3. Normal donors

Normal Donor bone marrows used for single cell RNA sequencing, flow cytometry analysis and T cell isolation for in vitro experiments were obtained from cryopreserved samples collected under an IRB approved protocol (#201103258) at Washington University.

| Normal Donor UPN | Age | Sex | Sample Type | Assays performed |
|---|---|---|---|---|
| ND050119 | 36 | female | Bone Marrow | Single Cell RNA sequencing and Flow Cytometry |
| ND050819 | 33 | male | Bone Marrow | Single Cell RNA sequencing, Flow Cytometry and T cell isolation for in vitro experiments |
| ND021920 | 40 | female | Bone Marrow | Flow Cytometry and T cell isolation for in vitro experiments |
| ND022620 | 25 | female | Bone Marrow | Flow Cytometry and T cell isolation for in vitro experiments |
| ND031820 | 33 | male | Bone Marrow | Flow Cytometry and T cell isolation for in vitro experiments |
| ND090617 | 25 | female | Bone Marrow | Flow Cytometry and T cell isolation for in vitro experiments |
| ND062619 | 38 | male | Bone Marrow | Flow Cytometry and T cell isolation for in vitro experiments |
| ND091119 | 26 | female | Bone Marrow | Flow Cytometry and T cell isolation for in vitro experiments |

**Table 1.** Normal donor samples used in this study.

#### C. AML Sample usage summary

| UPN | Source | Presentation: Exome sequencing | Remission: Error Corrected Sequencing | Bulk RNA-seq | sc-RNA seq | T cell activation experiments | LAG3 inhibition experiments |
|---|---|---|---|---|---|---|---|
| 139406 | Wash. U. | x | x | x | | x | x |
| 334228 | Wash. U. | x | x | x | x | | x |
| 627523 | Wash. U. | x | x | x | x | x | x |
| 759361 | Wash. U. | x | x | x | | x | x |
| 807970 | Wash. U. | x | x | x | x | | x |
| 854862 | Wash. U. | x | x | x | x | N/A | N/A |
| 868442 | Wash. U. | x | x | x | x | x | x |
| 964973 | Wash. U. | x | x | x | | x | x |
| 412761 | Wash. U. | x | N/A | x | | N/A | N/A |
| 635258 | Wash. U. | x | N/A | x | | x | x |
| 831711 | Wash. U. | x | N/A | x | | x | x |
| 976838 | Wash. U. | x | N/A | x | x | | x |
| 973536 | Wash. U. | x | N/A | x | | | x |
| 285 | Wash. U. | x | N/A | x | | N/A | N/A |
| 452442 | Wash. U. | x | N/A | x | | x | x |
| 916462 | Wash. U. | x | N/A | x | x | | x |
| 727185 | Wash. U. | x | N/A | x | | x | x |
| 613389 | Wash. U. | x | N/A | x | | x | x |
| 318748 | Wash. U. | x | N/A | x | x | N/A | N/A |
| 95-173-1 | Alliance | x | N/A | N/A | N/A | N/A | N/A |
| 97-0399-2 | Alliance | x | N/A | N/A | N/A | N/A | N/A |
| 00-0170-3 | Alliance | x | N/A | N/A | N/A | N/A | N/A |
| 04-0630-4 | Alliance | x | N/A | N/A | N/A | N/A | N/A |
| 06-0076-5 | Alliance | x | N/A | N/A | N/A | N/A | N/A |
| 08-0734-6 | Alliance | x | N/A | N/A | N/A | N/A | N/A |
| 08-3462-7 | Alliance | x | N/A | N/A | N/A | N/A | N/A |
| 09-1390-8 | Alliance | x | N/A | N/A | N/A | N/A | N/A |
| 10-2963-9 | Alliance | x | N/A | N/A | N/A | N/A | N/A |

**Table 2.** LFR sample studies. N/A= sample not available

| UPN | Source | Exome sequencing | Bulk RNA-seq | sc-RNA seq | T cell activation experiments | LAG3 inhibition experiments |
|---|---|---|---|---|---|---|
| 115225 | Wash. U. | x | x | | | x |
| 150288 | Wash. U. | x | x | | | x |
| 311636 | Wash. U. | x | x | | | x |
| 141273 | Wash. U. | x | x | | | x |

| | | | | | | |
|---|---|---|---|---|---|---|
| 452198 | Wash. U. | **x** | **x** | | **x** | **x** |
| 418499 | Wash. U. | **x** | **x** | | | **x** |
| 906708 | Wash. U. | **x** | **x** | | | **x** |
| 287 | Wash. U. | **x** | **x** | | N/A | |
| 104851 | Wash. U. | **x** | **x** | | | **x** |
| 161510 | Wash. U. | **x** | **x** | | **x** | **x** |
| 186706 | Wash. U. | **x** | **x** | | **x** | **x** |
| 208027 | Wash. U. | **x** | **x** | | N/A | |
| 296361 | Wash. U. | **x** | **x** | | | **x** |
| 329614 | Wash. U. | **x** | **x** | | **x** | **x** |
| 332131 | Wash. U. | **x** | **x** | **x** | **x** | **x** |
| 375182 | Wash. U. | **x** | **x** | | | **x** |
| 433325 | Wash. U. | **x** | **x** | | | **x** |
| 509754 | Wash. U. | **x** | **x** | | | **x** |
| 606061 | Wash. U. | **x** | **x** | | | **x** |
| 312340 | Wash. U. | **x** | **x** | | N/A | |
| 869922 | Wash. U. | **x** | **x** | **x** | **x** | **x** |
| 186481 | Wash. U. | **x** | **x** | | | **x** |
| 723101 | Wash. U. | **x** | **x** | | | **x** |
| 992966 | Wash. U. | **x** | **x** | | N/A | |
| 823477 | Wash. U. | **x** | **x** | **x** | N/A | |
| 508084 | Wash. U. | **x** | **x** | **x** | **x** | **x** |
| 869586 | Wash. U. | **x** | **x** | **x** | **x** | **x** |
| 721214 | Wash. U. | **x** | **x** | **x** | **x** | **x** |
| 875663 | Wash. U. | **x** | **x** | **x** | **x** | **x** |
| 126620 | Wash. U. | **x** | **x** | | | **x** |
| 286032 | Wash. U. | **x** | **x** | | | **x** |

**Table 3**. SFR sample studies . N/A= sample not available

### D. Statistical analyses
#### D1. Multivariate Analysis

**Supplemental Table 12. Multivariate Analysis of Factors Associated with CD4 Activation Status.***

| **Predictor** | **Odds Ratio (95% CI)** |
|---|---|
| Sex of patient | |
| Male | 0.74 (0.14–3.79) |
| Female (reference) | 1.00 |
| Age† | 0.99 (0.92–1.06) |
| WBC† | 0.98 (0.96–1.00) |
| BM blast percentage† | 0.99 (0.95–1.03) |
| ELN risk | |
| Favorable | 8.18 (0.75–89.65) |
| Intermediate or Adverse (reference) | 1.00 |
| NPM1‡ | 0.63 (0.03–12.92) |
| FLT3ITD‡ | 2.46 (0.15–39.83) |
| FLT3TKD‡ | 1.11 (0.06–19.08) |
| DNMT3A‡ | 0.46 (0.07–2.96) |
| NRAS‡ | 2.22 (0.34–14.45) |
| WT1‡ | 0.18 (0.01–3.55) |
| TET2‡ | 2.24 (0.18–28.31) |
| CEBPA‡ | 0.15 (0.01–2.48) |
| PTPN11‡ | 0.99 (0.03–28.37) |
| IDH2‡ | 3.45 (0.28–42.04) |
| IDH1‡ | 2.35 (0.18–30.51) |
| RUNX1‡ | 0.48 (0.03–7.82) |
| RAD21‡ | 3.17 (0.06–161.27) |
| SMC1A‡ | 1.87 (0.03–125.07) |
| SRSF2‡ | 8.07 (0.19–347.90) |

* N = 50. Tjur $R^2$ = .36. Deviance $\chi^2(20) = 26.49$, p = .15. Hosmer-Lemeshow $\chi^2(8) = 6.64$, p = .58.

† The odds ratio is for each 1-unit increase in the predictor.

‡ The odds ratio is for positive as compared to negative (reference).

Using the brglm2 0.8.0 package in R 4.0.2, a Firth logistic regression predicting CD4 activator status from all covariates of interest was fit to N = 50 observations (AMLs) using data contained in Supplementary table 11. The results of this multivariate analysis are summarized in Supplementary Table 12, where the odds ratios reflects the probability of CD4 activation. The model failed to identify any significant predictors of activator status. The model was not significantly better than the null model, Tjur $R^2$ = .36, $\chi^2(20) = 26.49$, p = .15, and a Hosmer-Lemeshow test suggested adequate fit, $\chi^2(8) = 6.64$, p = .58. Results of the multivariat e analysis are summarized in the table below.

##### D2. Additional statistical tests

Comparisons between the two groups were performed using Student t test (unpaired, 2-tailed) unless otherwise indicated in the main text. Statistical significance was accepted for p values <0.05.

P-values for multinomial tests were computed with 2-way ANOVA followed by post-hoc Tukey test using GraphPad Prism (software version 9.0.2), unless otherwise specified in the main text.

#### E. Sequencing and Bioinformatic Analyses

##### E1. Exome sequencing

DNA libraries were captured using either the SeqCap EZ Exome Probes v3.0 (Nimblegen/Roche), or the xGen Lockdown Exome Panel (IDT), supplemented with additional probes for recurrently mutated genes in AML. One case (831711) had whole genome sequencing as part of the TCGA AML study[1], but only variants within the exome space were used for this study. Cases with matched normal samples (skin or buccal swab) had somatic variants called as previously described[2]. Matched control DNA was not available for the Alliance cases, so variants were called using the pipeline described in (https://github.com/genome/analysis-workflows/blob/c73a2153e5a0fe04848f0f8e30093a11fa5b4712/definitions/pipelines/wgs.cwl). Briefly, variants detected with VarScan[3] were meshed with calls derived from GATK HaplotypeCaller[4] at cancer hotspots defined in DoCM[5] and filtered. Since somatic status is impossible to infer without a matched normal sample, mutation calls for these samples were restricted to those in protein-coding regions of known cancer-related genes that were predicted to result in amino-acid changes[1,6,7]. All variants were manually reviewed to remove sequencing artifacts[8]. Visualization was performed with the GenVisR[9] package for R and ggplot2[10].

##### E2. FLT3-ITD allelic ratio calculation

FLT3 allelic ratio (AR) was inferred from variance allele frequency (VAF) using the following formula:

AR = VAF / (1-VAF)

because AR = mutant allele VAF(s) / wild type allele VAF

If multiple FLT3 mutations were detected in one allele, the sum of every VAF(s) deteted for that allele was used in the formula.

AR ≥ 0.5 was considered as FLT3-ITD positive. AR < 0.5 was considered FLT3-ITD low.

#### E3. Error-Corrected Sequencing

A custom Agilent Haloplex panel was created, starting with the previously described Myeloseq panel[11] containing 40 recurrently mutated genes in AML. Sites of somatic mutations from previously sequenced exomes were added when possible, along with probes for *MYC* and several probes tiling SNPs in the *CDKN2A* region to detect loss of heterozygosity. The average coverage of all remission samples was 3042x (range 161-13,266x). Error correction was performed as described in Duncavage, et al[11]. For each sample, error-corrected readcounts were obtained at the site of every variant detected in the presentation sample. A site-specific error profile was created using samples lacking a given mutation as negative controls, and only variants significantly above the background level are retained, as in Xia, et al.[12]

#### E4. Total RNA sequencing from unfractionated bone marrow samples

Total RNA sequencing was performed as described previously[1]. Gene expression was quantified with kallisto[13] version 0.43.1 using human transcripts from Ensembl version 95. and differentially expressed genes (DEGs) detected using edgeR[14]. Genes expressed at very low levels (CPM < 1 in over 50% of the samples) were excluded. After initial analyses found no differentially expressed genes between SFR and LFR groups, the observation was made that expression heterogeneity driven by AML FAB subtype was a confounding variable. Four groups were included in a subsequent design: FAB types M1, M2, M4, and M5. After applying multiple testing correction, no DEGs were found to be significantly different between LFR and SFR even after using FAB as a covariate.

#### E5. Single Cell RNA sequencing

##### E5.1. Cell Preparation and Sorting

AML cells were thawed from cryovials into 13 ml of 100% FBS and immediately concentrated by centrifugation. The pellet was resuspended in 1 to 5 ml of PBS supplemented with 2% FBS for cell counts. 1 ul/ml of Sytox Red was added to the cell suspension and incubated at room temperature for 15 min. Cells were then filtered through a 70 uM screen to remove any clumps prior to cell sorting for Sytox negative cells (i.e. viable cells). Sorting was performed on a Synergy SY3200 BSC cell sorter (Sony Biotechnology, San Jose, CA). Subsequent to sorting, cells were centrifuged and resuspended in 1-3 ml of PBS +0.04% BSA for cell counts. One million cells were transferred to one Eppendorf tube at a concentration of 1,000 cells/uL, and submitted for single cell sequencing. cDNA libraries were prepared from individual cells using the 10x Genomics Chromium Single Cell 5′ Kit as described in Petti, et al.[15]

##### E5.2. Single Cell Analyses

Sequence was generated with 151 bp reads on the Illumina NovaSeq platform. Alignments were produced using Cellranger version 3.1.0 and the provided annotation package based on Ensembl version 93 (https://support.10xgenomics.com/single-cell-gene-expression/software/overview/welcome). Sequencing saturation was >90% for all samples, indicating adequate transcript sampling. Single cells were assigned to lineages using cellMatch[16], and expressed mutations were detected using cbSniffer (https://github.com/sridnona/cb_sniffer). Gene/barcode matrices, lineage assignments, and mutation/barcode lists were imported to the Partek Flow software (build version 10.0.20.1231), where all subsequent analyses were performed. QA/QC filtering was performed according to the following parameters: total read counts greater than 500, mitochondrial reads <13% and expressed genes >500. 91473 of 123605 cells (74%) passed the QA/QC filter. Counts were normalized as recommended (log transformation, converted to counts per million and +1).

Principal component analysis (PCA) was performed prior to graph- based clustering or UMAP visualizations. Genetically-defined AML cells were classified as previously described[15], selected and filtered for independent PCA analysis, clustering and downstream statistical analysis.

Differential gene expression analysis for AML cells was performed using ANOVA with significance thresholds of FDR<0.001 and fold changes +/- 2. Genes with fold changes >1.5 were selected for Gene Ontology analyses. Only pathways with >10 genes and with FDR<0.05 were considered as significantly enriched. Targeted analyses on genetically-defined AML cells to identify cells with an active ‘gene set’ (i.e. gene signatures such as DNA repair genes) were performed with the AUCell method. AUCell uses the “Area Under the Curve” (AUC) to calculate whether a critical subset of the input gene set is enriched within the expressed genes for each cell. The distribution of AUC scores across all the cells permit exploration of the relative expression of the signature. Since the scoring method is ranking-based, AUCell is independent of the gene expression units and the normalization procedure.

The analyses of immune cells were performed similarly. Graph-based clustering was used to identify subsets of immune cells, then biomarkers for each cluster were calculated using an ANOVA test where each cluster is compared to the other cells in the data set. Genes with fold-change > 1.5 were included, and sorted by ascending p-value (ties broken by greater fold change). The gene lists are available in Supplementary Table 7. Top marker genes for each cluster were evaluated with gene list enrichment analysis and candidate prioritization tools (https://toppgene.cchmc.org/enrichment.jsp) for cell type identification. Clusters belonging to the same cell type (i.e. all clusters of CD4 Naïve T cells) were pooled for graphical representation in Figure 3B. Because the transcriptional changes in non transformed immune cells were more subtle, the thresholds for significance of the ANOVA targeted analysis for the activation/exhaustion markers were set at FDR<0.05 and log fold changes of +/- 1.3.

#### E6. Digital Droplet PCR

All Digital Droplet PCR reagents were purchased from Bio-Rad. Assays with primers and probes for ddPCR were commercially available and designed by Bio-Rad as per MIQE guidelines[17]. Droplet generation was obtained with the Bio-Rad QX100 and QX200 Droplet Digital PCR Systems and ddPCR was performed and analyzed as previously described[18]. Briefly, 200 ng of DNA was analyzed in each well, for a total of 1 ug of DNA across 5 wells per sample. DNA was digested with HindIII at 1uL/well to ensure maximum DNA accessibility. To optimize the annealing temperature, we first ran a plate with a temperature gradient to select the temperature that allowed for optimal droplet separation. Each well contained a 1:1 mixture of positive control DNA (the DNA containing the mutation, obtained from IDT) and the WT DNA (negative control, obtained from a normal donor) along with the specific assay for each mutation. Control wells contained WT DNA only (negative), G-block DNA only (positive), or no-template controls (blank). To determine the exact level of sensitivity or each assay, serial dilutions of each Mutant DNA into WT DNA were performed, and the Limit of Detection was calculated with the following formulas:

Limit of blank (LoB) = mean blank + 1.645(SD blank)

Limit of Detection = LoB + 1.645(SD low concentration of target)

The data was visualized and analyzed using Quanta Software (BioRad).

##### E7. Immunologic Analyses

MHC Class I and II typing was performed using Optitype[19] and PHLAT[20], both with default parameters. When more than two alleles were reported for a group (e.g. HLA-A), the two alleles with the highest scores were retained. Samples from the Alliance without matched normals or RNA-seq were not included in this analysis. HLA typing failed for one sample (318748) and it was therefore excluded. Neoantigen predictions were produced with pVACtools 2.0 alpha v8, using default parameters and all supported binding prediction algorithms[21]. Expressed candidate neoantigens were defined as mutations with at least one mutant peptide with median predicted binding (ic50) < 500nM and evidence that the mutation was expressed in the RNA (RNA VAF > 0). No significant differences were observed in neoantigen burden or in HLA types (Supplemental Figure 8). Differences between groups were assessed with a two-sided t-test in R version 4.0.2.

##### E8. Data Deposition

All sequence data produced for this study is deposited in dbGaP study phs000159 v.11.

#### F. In vitro Assays

##### F1. Culture of primary human AML cells

Cryovials of primary AML cells were quickly thawed into 10 mL of FBS (BioTechne) supplemented with 2 mM EDTA, and then centrifuged at 520 x G for 5 minutes. Cell pellets were resuspended at 0.5-1 x $10^6$/ mL in R10 media: RPMI supplemented with 10% FBS, 1% penicillin/streptomycin (GibCo), 1% GlutaMax (GibCo), 1% non-essential amino acids (GibCo), 1% sodium pyruvate (Corning), 2% HEPES (Corning), 900uL/L β-mercaptoethanol, and human cytokines (PeproTech) including: SCF (100ng/mL), FLT3 (10ng/mL), TPO (10ng/mL), IL3 (10ng/mL), IL6 (20ng/mL), IL2 (10ng/mL), and used for in vitro experiments.

#### F2. T cell cultures

Cryopreserved primary AML samples were rapidly thawed into 10 mL FBS (Biotechne) and centrifuged at 520 G for 5 minutes. Cell pellets were resuspended in PBS (Hyclone) supplemented with 0.5% BSA and 2 mM EDTA, and stained using Miltenyi Pan-T Isolation Kit according to manufacturer's instructions. Cells were separated using autoMACS Pro Separator, and the non-labelled fraction (containing enriched CD3+ T-cells) was resuspended in R10 media, described in Section F1.

#### F3. T cell stimulation assay

Cryopreserved bone marrow from healthy donors were thawed as described above. For T cell cultures in presence of AML cells, the percentage of CD3+ and CD45+ cells were determined with flow cytometry on a small aliquot of the sample prior to plating, and the absolute number of T cells within each sample was calculated by multiplying the percent CD3+ by the total number of viable cells. For T cell cultures in the absence of AML cells, CD3+ cells were separated as described in B 4.2, and CD3+ cells were counted after separation. T-cells were supplemented with a 1:1 ratio of GibCo Dynabeads Human T-Activator CD3/CD28 (Lot# 00847442) for 5 days prior to flow cytometry analysis.

#### F4. LAG3 inhibition assays

Each cryopreserved AML sample was thawed into 10 mL FBS (Biotechne), centrifuged at 520 x G for 5 minutes, resuspended in R10 media, and split equally into 3 wells of a 24-well plate. The percentage of CD3+ and CD45+ cells were determined with flow cytometry on a small aliquot of the sample prior to plating. Each well was treated with: 1) no added reagents, 2) GibCo Dynabeads Human T-Activator CD3/CD28 (Lot# 00847442) at a 1:1 ratio with T cells and mouse IgG1 isotype control (invitrogen, Lot# UG287713) at 1 ug/mL, or 3) T-Cell stimulation beads, and 1 ug/mL mouse anti-human LAG3 antibody (antibodies-online.com; clone 17B4). Cells were cultured for five days, and then counted prior to analysis using a Bio-Rad Yeti ZE5 flow cytometer.

### G. Flow cytometry Analyses

#### G1. T cell immunophenotyping

Immune profiling of AML samples was performed on a Bio-Rad Yeti ZE5 using a modified protocol developed by Thermo Fischer[22]. Cells were stained with the following combination of antibodies to identify indicated cell types: FITC conjugated anti-human CD134/OX40 (Invitrogen; clone ACT 35), PerCP-Cy5.5 conjugated anti-human CD45RA (Invitrogen; clone HI100), eFluor 660 conjugated anti-human CD183/CXCR3 (Invitrogen; clone CEW33D), AF700 conjugated anti-human CD196/CCR6 (Invitrogen; clone R6H1), BV421 conjugated anti-human CD8 (Biolegend; clone RPA-T8), BV510 conjugated anti-human CD127 (Biolegend; clone A019D5), Superbright 600 conjugated anti-human CD4 (Invitrogen; clone SK3), BV650 conjugated anti-human CD25 (Biolegend; clone BC96), BV711 conjugated anti-human CD3 (Biolegend; clone OKT3), BV786 conjugated anti-human CD279/PF-1 (BD Biosciences; clone EH12.1), PE conjugated anti-human CD278/ICOS (Invitrogen; clone ISA-3), PE-eF610 conjugated anti-human CD223/LAG3 (Invitrogen; clone 3DS223H), Pe-Cy7 conjugated anti-human CD194/CCR4 (BD Biosciences; clone 1G1). Viability was assessed with LIVE/DEAD™ Fixable Near-IR Dead Cell Stain Kit (ThermoFisher, cat # L10119).

FMO controls were used for gate placement and files were analyzed with FlowJo (version 10.7.1). An example of gating and population hierarchy is provided below.

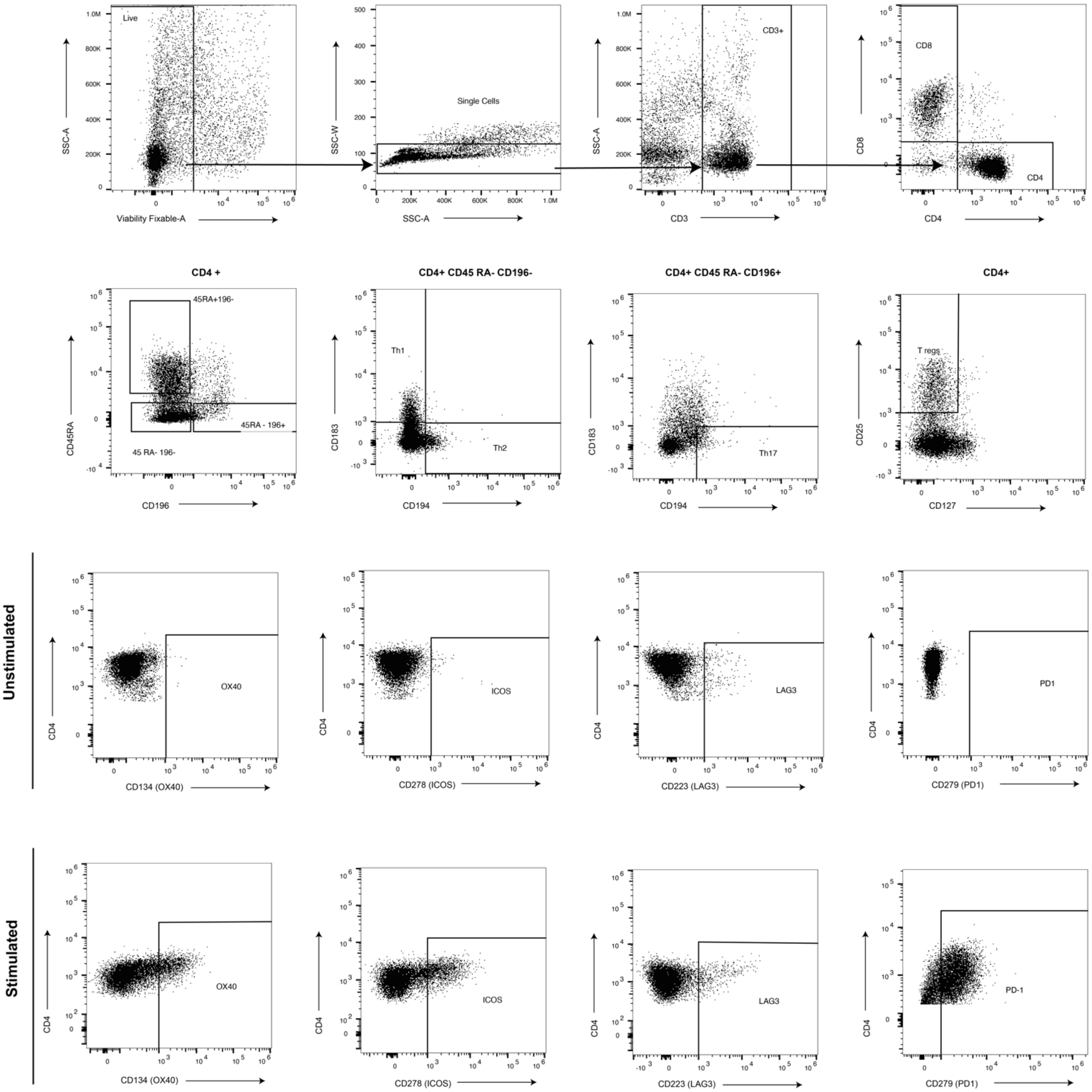

Live
SSC-A
Viability Fixable-A
Single Cells
SSC-W
SSC-A
CD3+
SSC-A
CD3
CD8
CD4
CD8
CD4
CD4 +
45RA+196-
45RA - 196+
45 RA- 196-
CD45RA
CD196
CD4+ CD45 RA- CD196-
Th1
Th2
CD183
CD194
CD4+ CD45 RA- CD196+
Th17
CD183
CD194
CD4+
T regs
CD25
CD127
Unstimulated
OX40
CD4
CD134 (OX40)
ICOS
CD4
CD278 (ICOS)
LAG3
CD4
CD223 (LAG3)
PD1
CD4
CD279 (PD1)
Stimulated
OX40
CD4
CD134 (OX40)
ICOS
CD4
CD278 (ICOS)
LAG3
CD4
CD223 (LAG3)
PD-1
CD4
CD279 (PD1)

### H. Supplementary Figures

#### Supplementary Figure 1

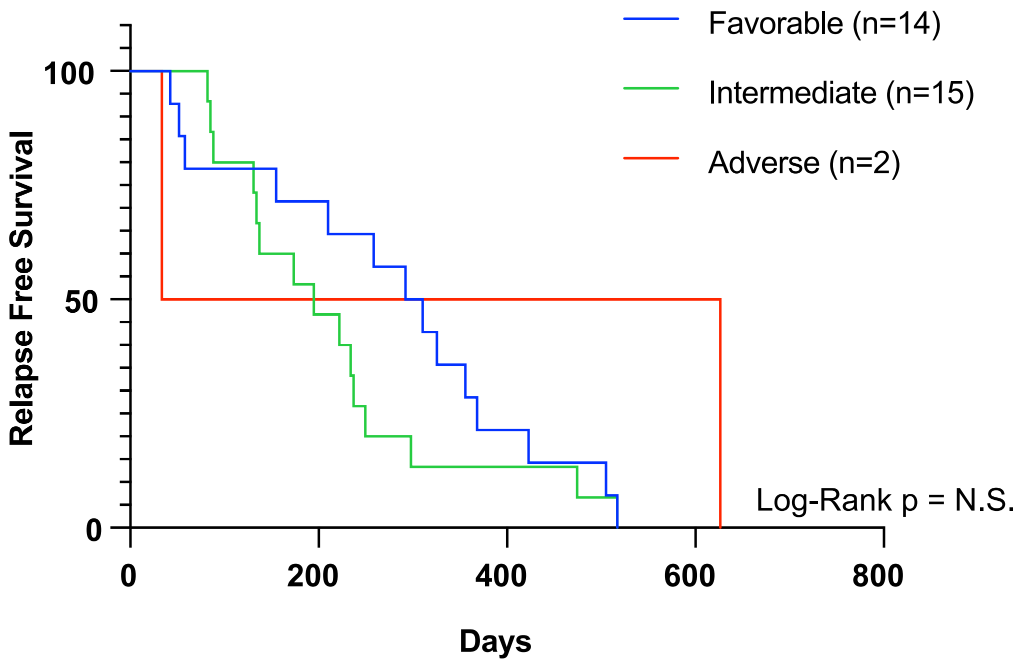

**Supplementary Figure 1. Relapse free survival curve of SFR patients by ELN risk category.** Kaplan Meier curves illustrating Relapse Free Survival of SFR patients. Curves are stratified based on ELN risk category, as indicated in the legend. No statistical differences in survival between groups were detected using log rank method.

#### Supplementary Figure 2

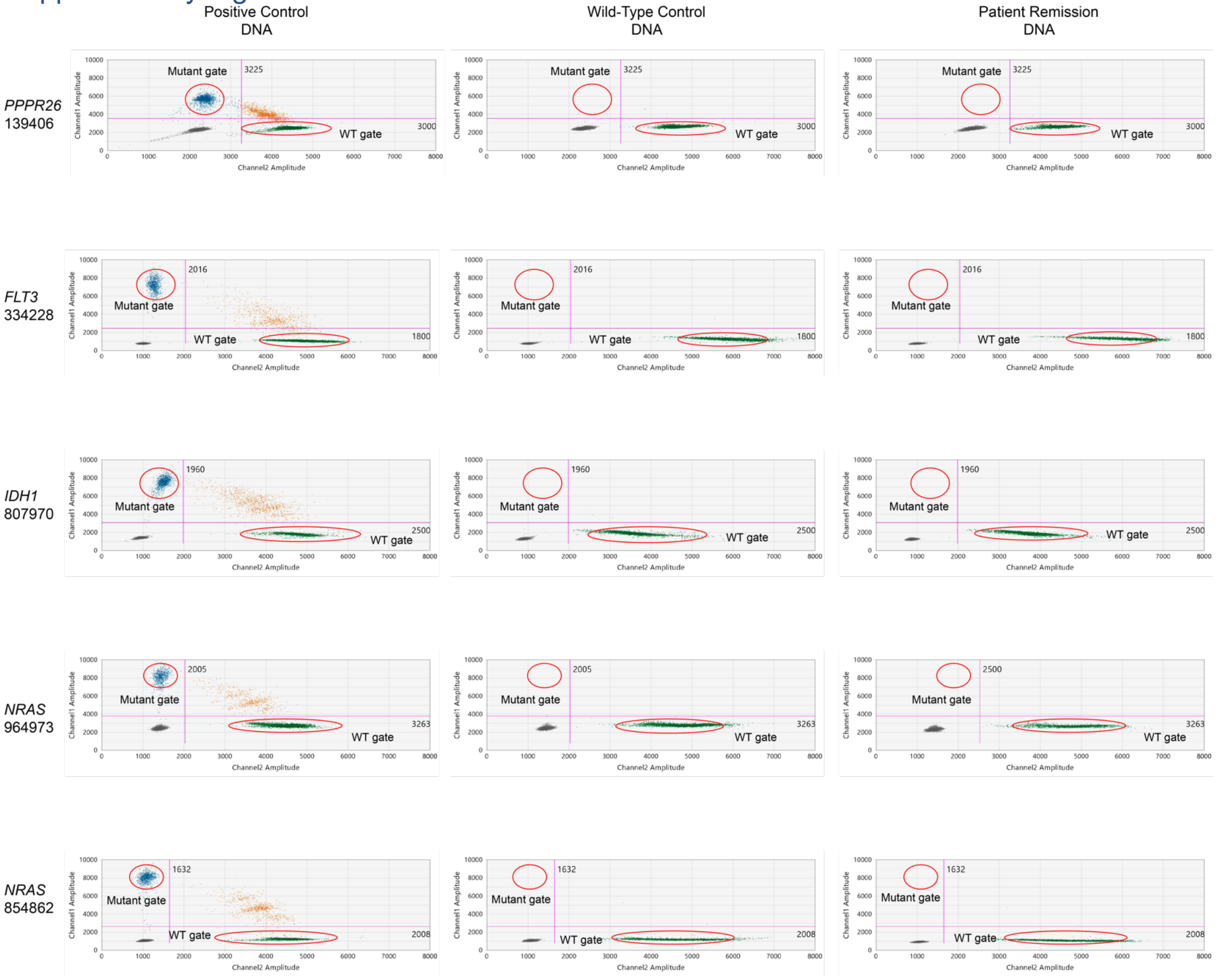

**Supplementary Figure 2. Digital droplet PCR evaluation of selected founding clone mutations in 5 LFR cases in remission.** Digital droplet PCR scatter plots showing the clearance of a selected point mutation (one in each row) in 5 long first remission cases that contained a founding clone mutation for which there were commercial reagents available from Bio-Rad. The analysis was performed using the longest remission sample for each patient. Droplets containing only the mutant allele are blue, droplets containing both the wild type and the mutant allele are orange, droplets containing only the wild type allele are green, and empty droplets are gray. The first two columns show positive controls (G-blocks), and negative controls (DNA form a normal donor), respectively. The last column represents the results in the patient DNA from the remission sample. This Digital Droplet PCR assay was employed to increase the sensitivity of mutation detection in the remission samples to 1 cell in 100,000 cells (0.001%). The sensitivity was estimated based on limit of detection calculations as detailed in Supplementary Methods.

#### Supplementary Figure 3

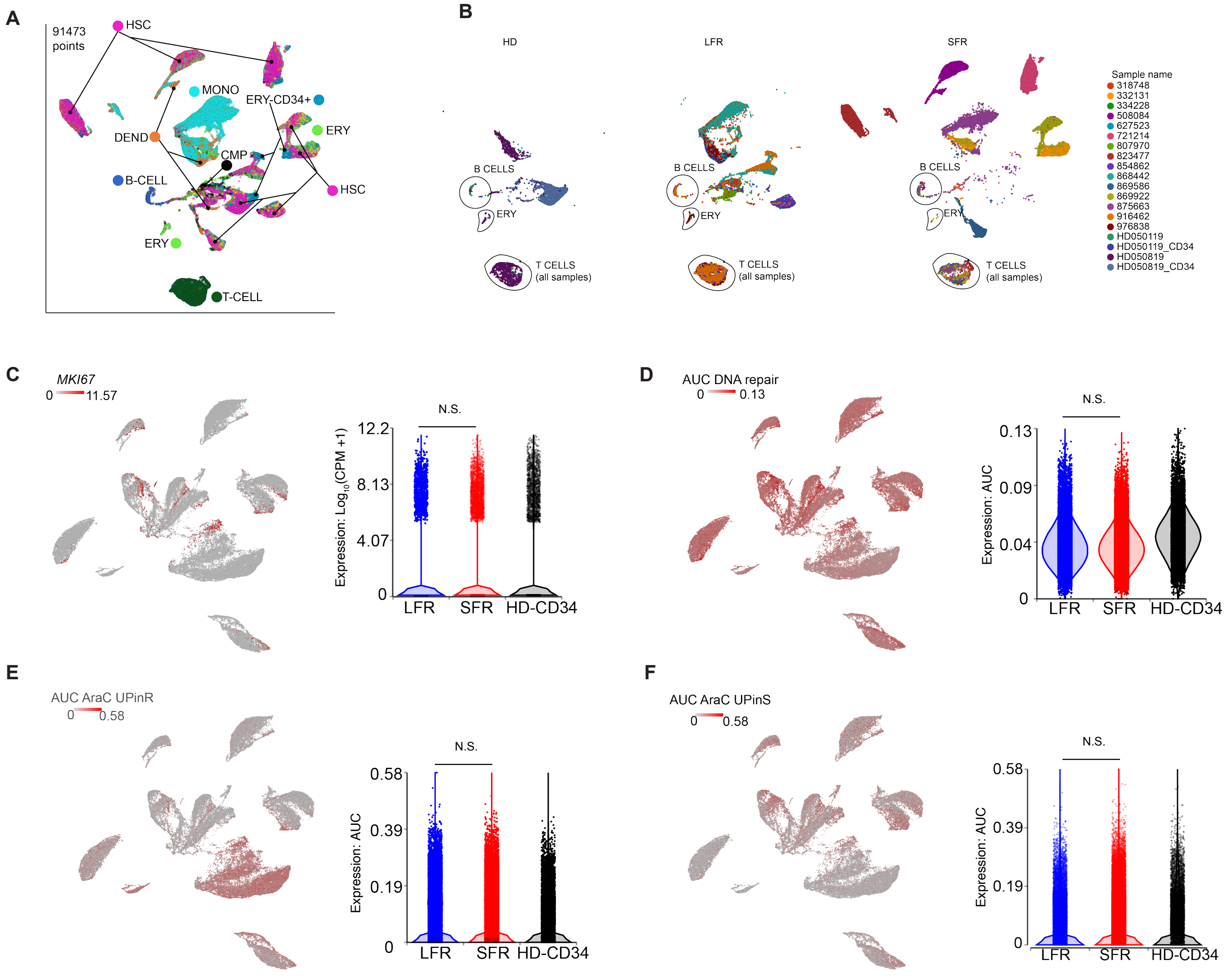

**Supplementary Figure 3. Single cell RNA-sequencing (scRNA-seq) analyses of defined chemotherapy response pathways. Panels A** and **B.** UMAP projections of scRNA-seq data from 8 LFR patients, 7 SFR patients and 2 healthy donor samples (the latter represents both unfractionated BM and CD34+ sorted cells from the same donors). **Panel A.** All cells from all samples are colored by lineage based on CellMatch[16]. **Panel B.** Cells are split by group (HD, LFR and SFR) and colored by sample ID. Black lines outline clusters containing cells from all samples, the majority of which are T cells, along with smaller clusters of B cells and erythroid cells.

**Panels C through F.** UMAP projections of scRNA-seq data from genetically-defined AML cells and from sorted healthy donor BM-derived CD34+ cells (same projection depicted in Figure 2B), where each cell is colored according to the levels of expression of: **Panel C**) the proliferative marker *MKI67* (average expression),

**Panel D**) a DNA repair signature gene list (AUC cell, full list in **Supplementary Table 4**), **Panel E**. Genes associated with AraC resistance (*CDA, DCTD, SAMHD1, NT5C2, RRM1, RRM2,* AUC cells). **Panel F**. Genes associated with AraC sensitivity (*SLC28A3, SLC29A1, DCK, CMPK1, NME1;* AUC cells). Area under the curve (AUC) is a rank-based scoring method that is independent of sample size, gene expression unit, and normalization method (further details provided in Supplementary Appendix Section E4.2). The color gradient is from grey (no expression) to red (high expression). N.S.= not significant.

#### Supplementary Figure 4

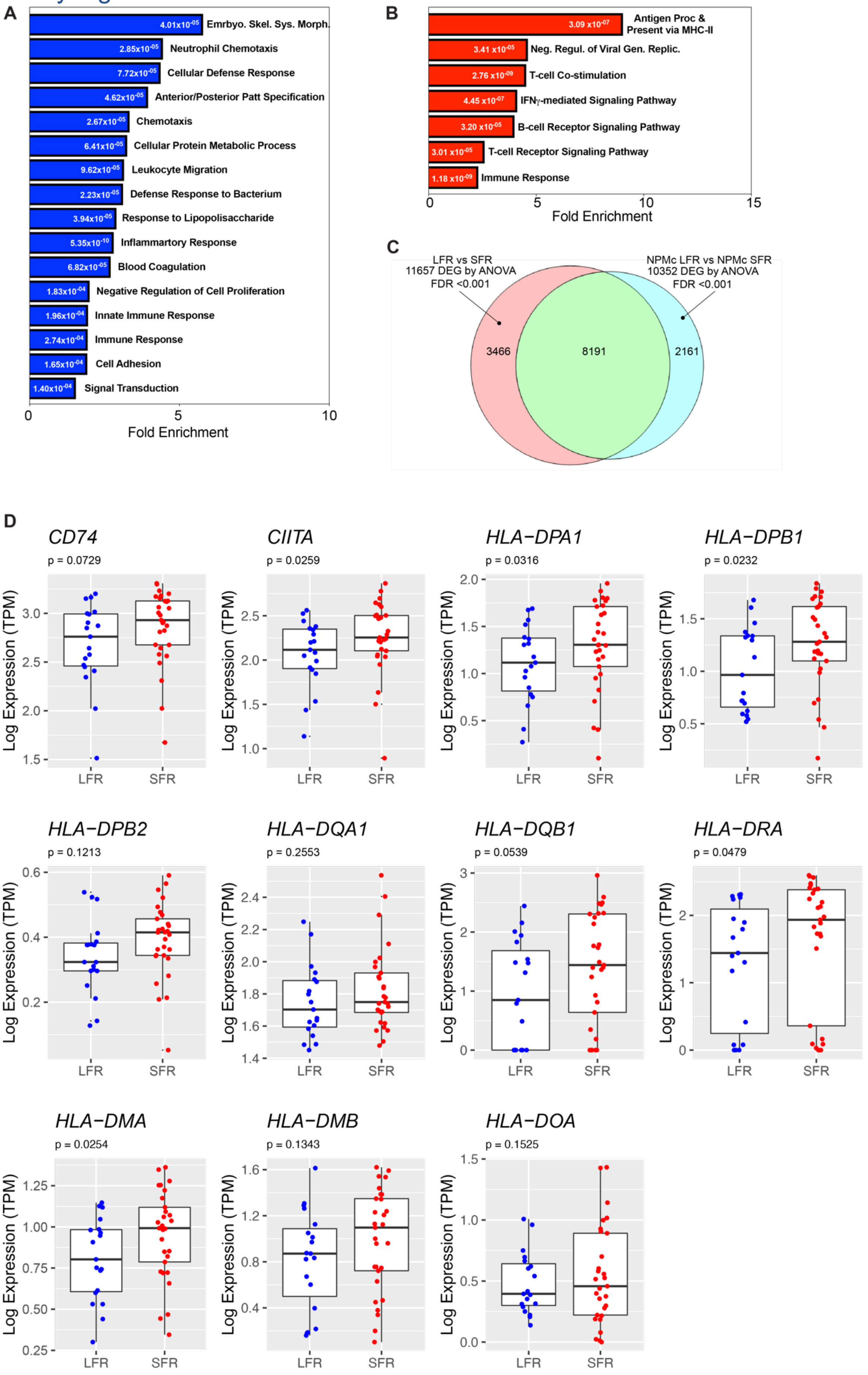

**Supplementary Figure 4. Enrichment of adaptive immune response pathways and HLA class II gene expression in SFR vs. LFR AMLs. Panels A and B.** Enriched Gene Ontology pathways of differentially expressed genes between LFR and SFR groups. Blue and red bars are pathways enriched in the LFR or SFR cases, respectively. Numeric values indicate the FDR for each specific pathway. Each bar length is proportional to the fold enrichment for that pathway. **Panel C**: Venn diagram showing the relationships between 2 differentially expressed gene sets (as defined by ANOVA, FC $\pm$ 2 and FDR<0.001) obtained comparing LFR vs SFR (red circle) and *NPM1**LFR vs *NPM1**SFR (blue circle). 8191 genes were shared between the 2 comparisons (green overlapping area). **Panel D**. Boxplots showing the expression of *CIITA* (the master regulator of HLA class II expression), all the HLA class II genes that were detected in the LFR cases (blue dots) vs. SFR cases (red dots), and *CD74*, a gene that is also regulated by *CIITA*. The expression of 5/11 genes was significantly (p<0.05) higher in the SFR samples, while all genes showed a trend towards higher expression in the SFR samples.

#### Supplementary Figure 5

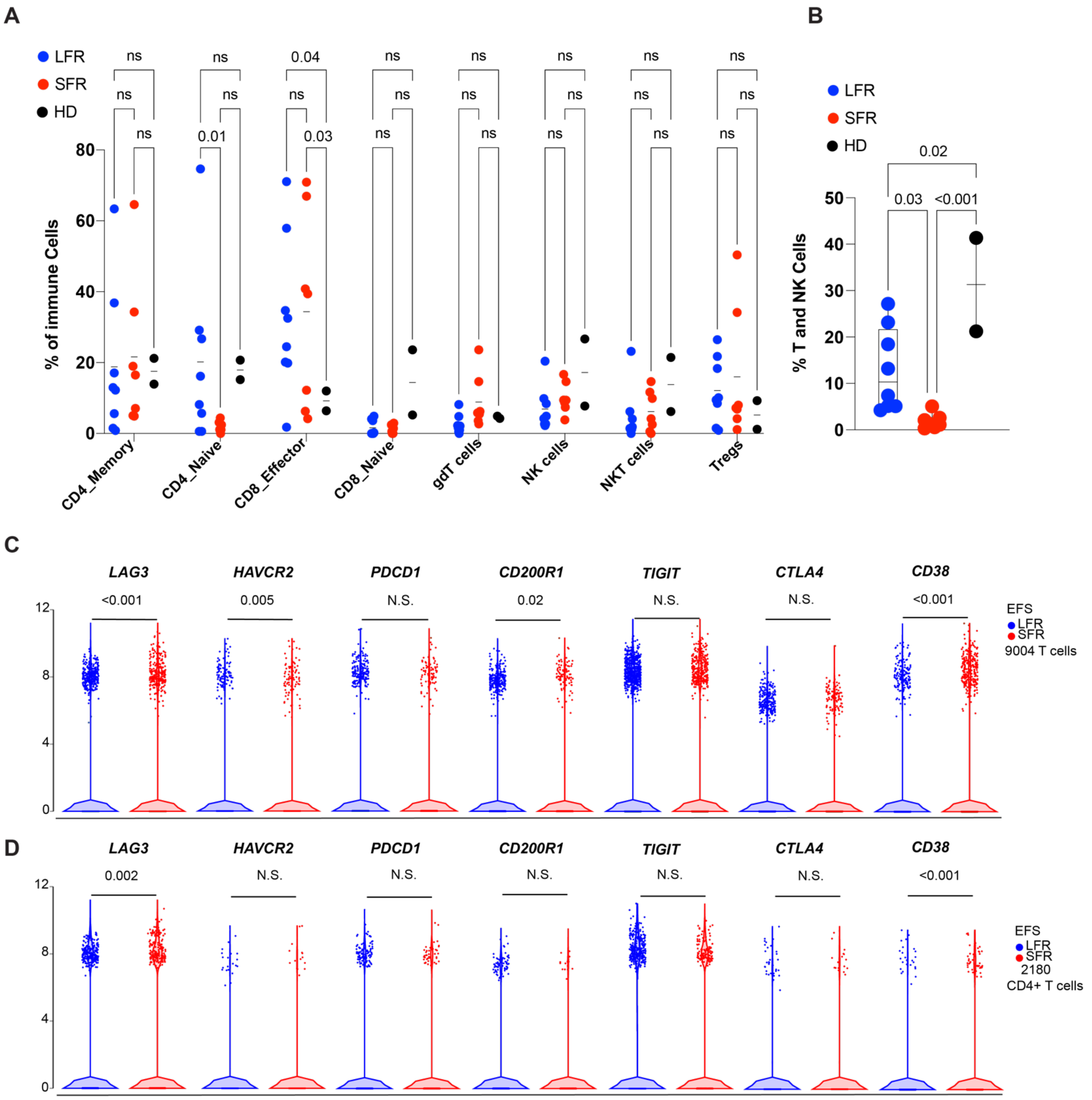

**Supplementary Figure 5. Differences in bone marrow T cell subsets and immune checkpoint gene expression in LFR vs. SFR patients, from scRNA-seq data. Panel A.** Scatterplots indicating the proportion of T cells in subsets defined by scRNA-seq in the presentation bone marrow samples, as defined by graph based clustering. Black line indicates the median value for each parameter tested; 2-way ANOVA and Tukey multiple comparison tests were used to test for significant differences between groups. **Panel B.** Boxplots indicating the proportion of T and NK cells as defined by lineage assignment in the scRNA-seq data; 2-way ANOVA and Tukey multiple comparison tests were used to test for significant differences between groups. **Panels C and D.** Violin plots showing the expression differences of defined immune checkpoint genes in 9,004 T and NK cells identified in the samples (**Panel C**, Supplementary table 9), and in the 2,180 CD4+ defined T cells (**Panel D** and Supplementary table 10). N.S.= not significant.

#### Supplementary Figure 6

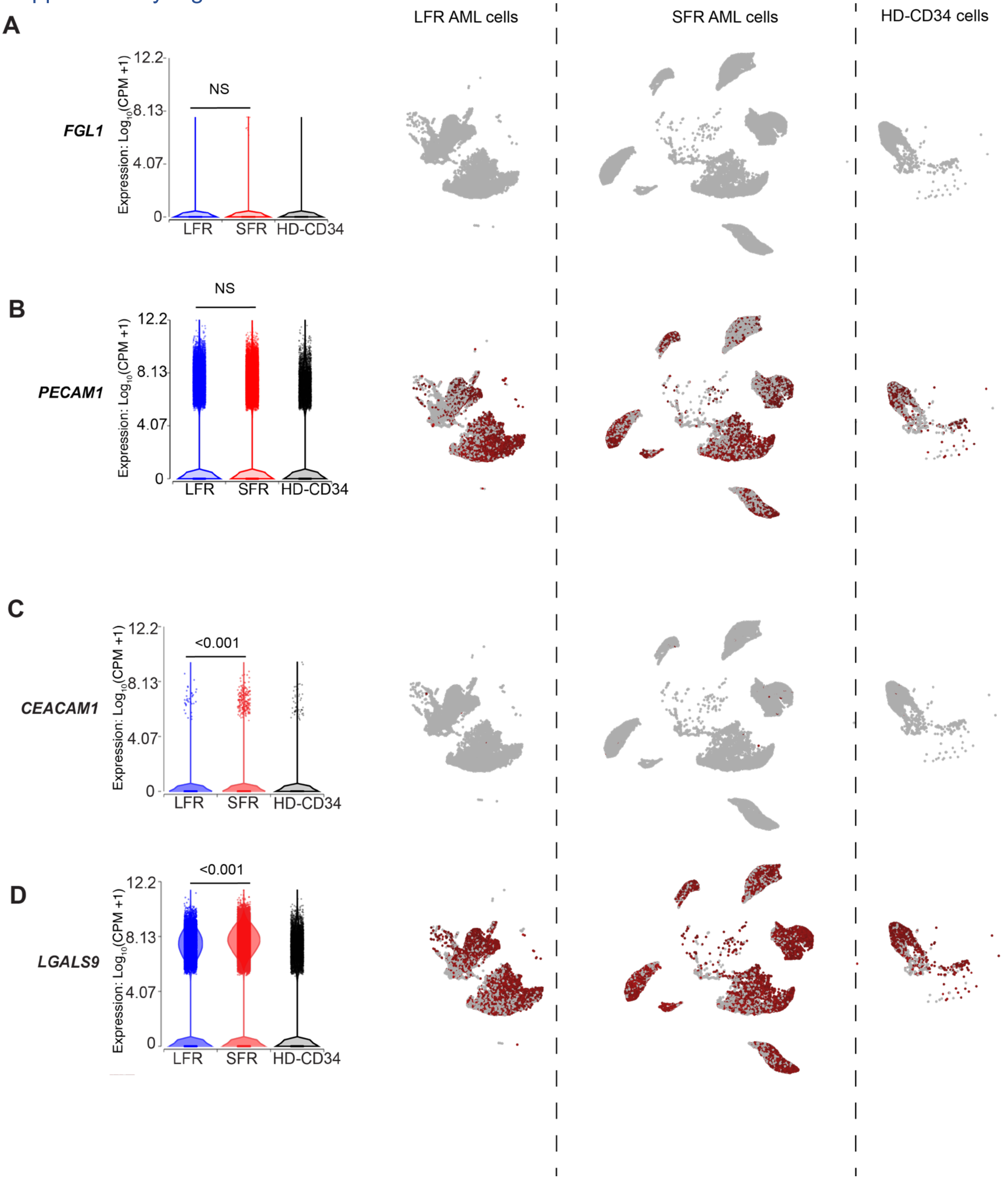

**Supplementary Figure 6. Expression of genes encoding immune checkpoint ligands in genetically-defined AML cells from LFR vs. SFR patients. Panels A through D:** Violin plots and relative UMAP projections split by response (LFR, SFR, HD) showing the expression levels of genes encoding ligands for checkpoint inhibitors in the genetically-defined AML cells of of LFR vs. SFR patients: **Panel A**. *FGL1*, a proposed alternative ligand for LAG3; **Panel B**. *PECAM1*, a ligand for CD38; **Panel C**. *CEACAM1* and **Panel D**. *LGALS9,* both ligands for *HAVCR2* (the gene encoding TIM3). Expression differences were determined using ANOVA comparisons from the scRNA-seq data, and p values are indicated above each violin plot. N.S.= not significant.

#### Supplementary Figure 7

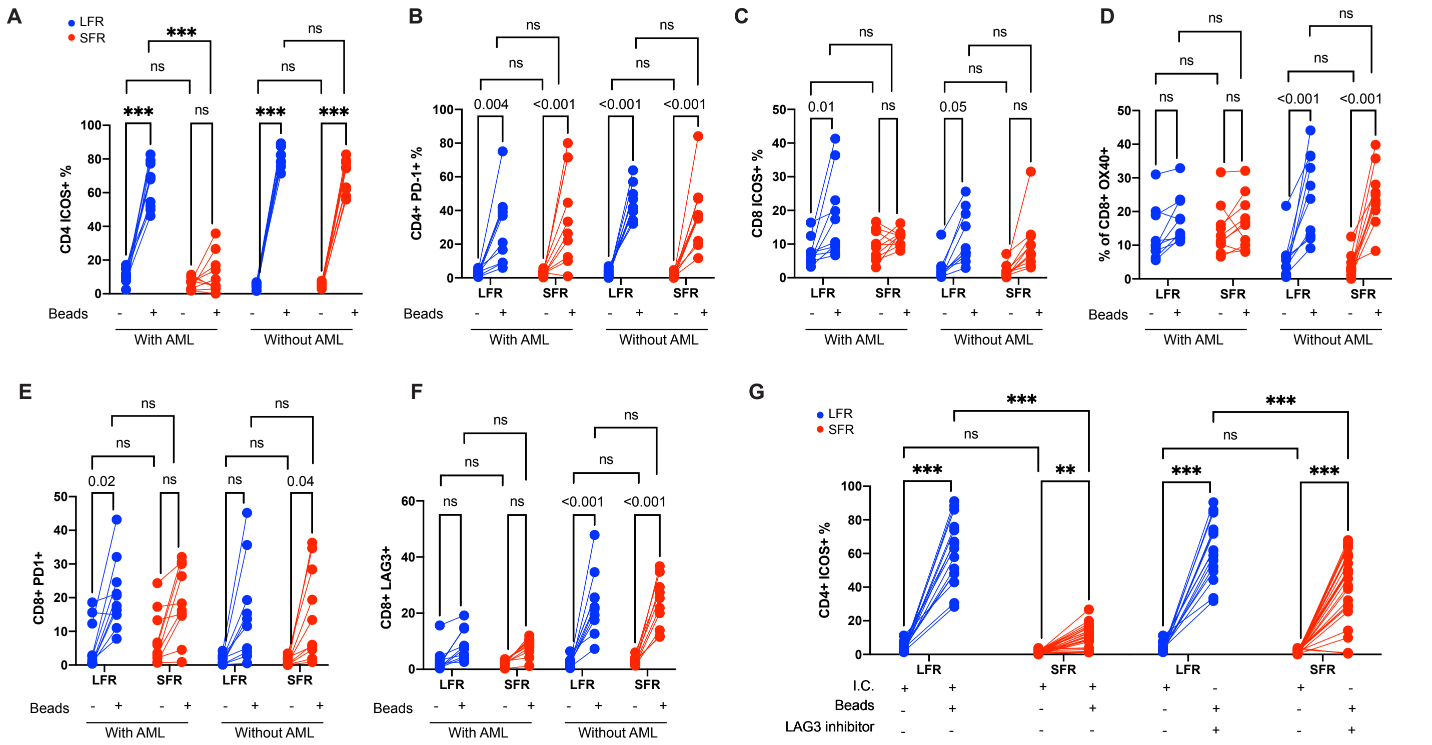

**Supplementary Figure 7. T cell responses to CD3/CD28 bead stimulation in the presence or absence of AML cells and T cell activation changes after LAG3 inhibition.**

**Panels A through E**. Line plots showing changes of surface ICOS (**Panel A**), PD-1 (**Panel B**) expression on CD4+ T cells; and ICOS (**Panel C**), OX40 (**Panel D**), PD1 (**Panel E**), LAG3 (**Panel F**) in CD8+ T cells in unstimulated and CD3/28 bead-stimulated endogenous T cells from unfractionated ("with AML") or CD3+ selected ("without AML") bone marrow samples from presentation. Results are the summary of 3 independent experiments (LFR, blue lines, n=10, SFR, red lines, n=10). 2-way ANOVA with Tukey correction for multiple comparison was used to calculate the p values shown in the graphs. Ns= not significant.

**Panel G** shows that similar levels of activation, as defined by ICOS expression, can be achieved in CD4+ T cells from SFR samples in the presence of a LAG3 blocking antibody at day 5 post stimulation. The negative controls were treated with an isotype matched antibody (I.C.= Isotype Control). 2-way ANOVA and Tukey multiple comparison tests were used to test for significance differences between groups. (n=15 for LFR and n=26 for SFR cases).

Supplementary Figure 8

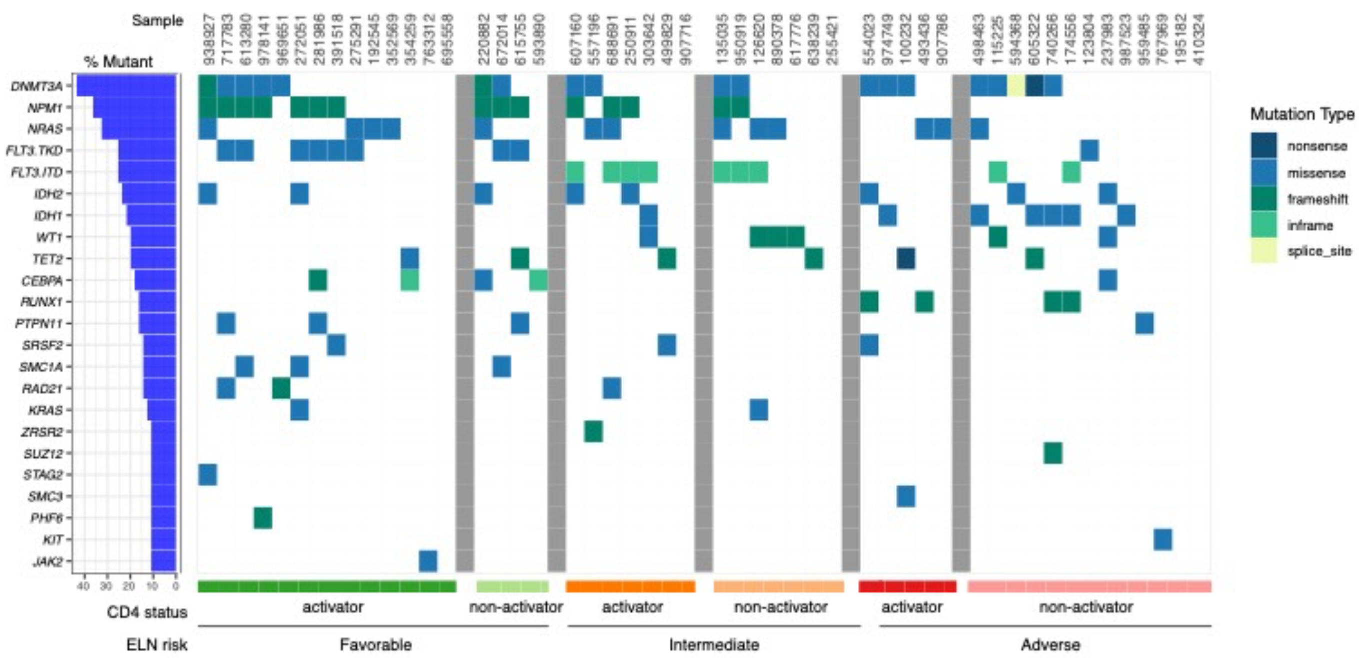

**Supplementary Figure 8. Mutational landscape of the 50 Extension cases.**
Each column is a case labeled by UPN, and each row represents a recurrently mutated AML gene. Mutations are ranked by frequency, as indicated by the blue bars to the left of the plot. The samples are grouped according to ELN categories and within each category, CD4 cell activator vs. non activator samples are separated by the grey vertical lines.

Supplementary Figure 9

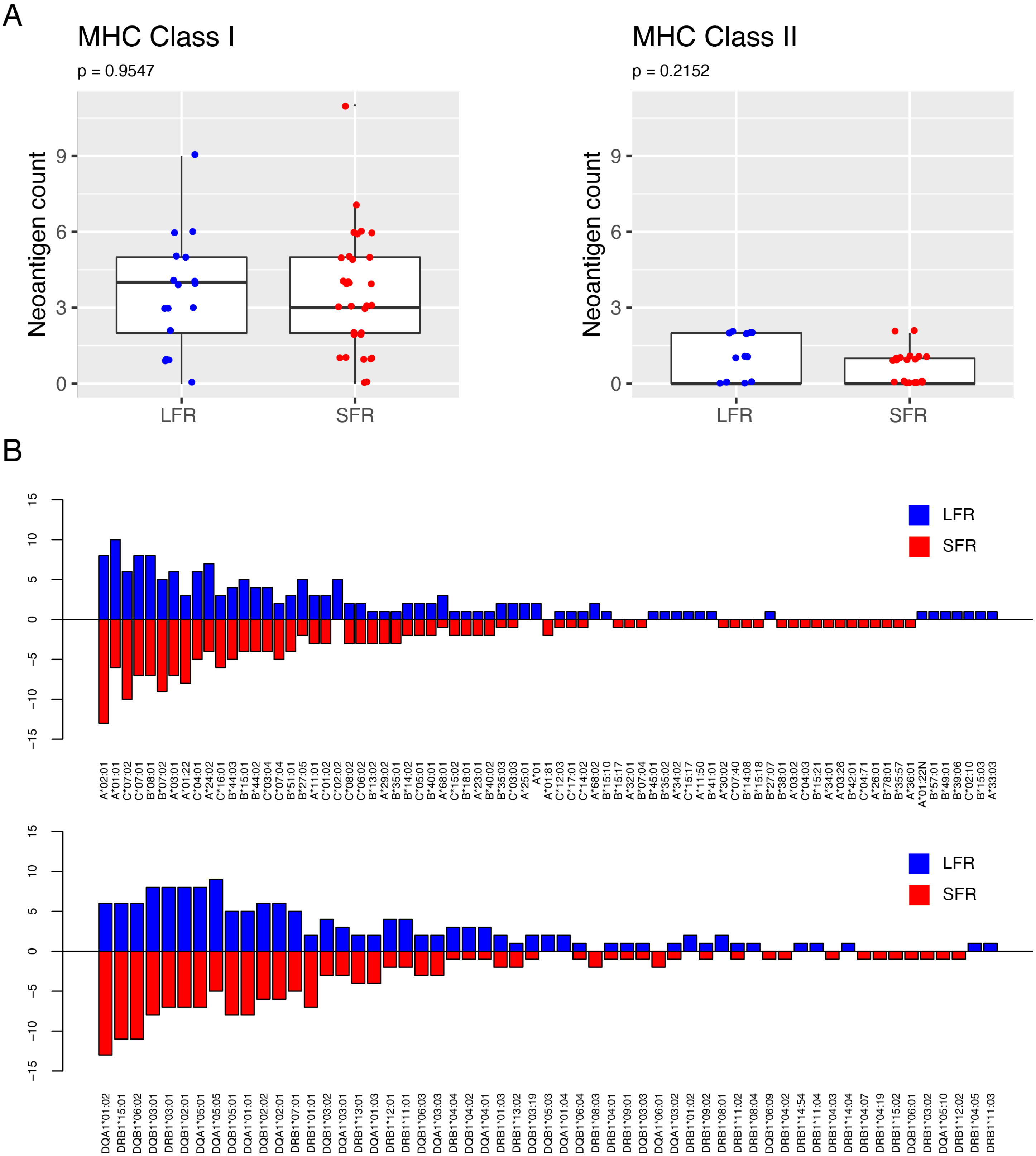

**Supplementary Figure 9. Evaluation of neoantigen load and HLA haplotypes in LFR vs. SFR patients.**

**Panel A.** Summary boxplots of total neoantigens predicted in the context of MHC class I (left panels) or MHC class II (right panels) for LFR (blue) and SFR (red) samples. **Panel B**. The proportion of LFR and SFR with a particular HLA type. There is no significant relationship between outcome and any specific HLA type observed in these cases.

### I. List of Supplementary tables
(see excel files uploaded separately)

##### Supplementary Table 1. LFR and SFR clinical characteristics
This table contains additional clinical information regarding each patient and the somatic mutations detected in each case.

##### Supplementary Table 2. LFR and SFR somatic mutations
This table lists all the somatic mutations discovered by exome sequencing in the 28 LFR and 31 SFR cases at diagnosis.

##### Supplementary Table 3. Haloplex results
This table contains the error-corrected sequencing results for the presentation and remission samples of selected LFR patients with remission samples available for analysis.

##### Supplementary Table 4. DNA repair gene list
This table lists the DNA repair genes used for the AUCell analysis shown in Supplementary figure 3D.

##### Supplementary Table 5. Single cell RNA sequencing results
This table shows the results of the ANOVA comparison between LFR and SFR samples obtained with single cell RNA sequencing data, and the relevant Gene Ontology analysis results.

##### Supplementary Table 6. Single cell RNA sequencing analysis results (NPM1 cases only)
This table shows the results of the ANOVA comparison between LFR and SFR samples with NPM1 mutations, using single cell RNA sequencing data and the relevant Gene Ontology analysis results.

##### Supplementary Table 7. List of shared DEGs between comparisons in S5 and S6
This table lists the shared DEGs between the lists generated from the comparisons in Tables S5 and S6, and the relevant Gene Ontology analysis results.

##### Supplementary Table 8. T cell graph-based clustering
This file shows the top marker genes in each cluster of T cells, ranked by p-value.

##### Supplementary Table 9. Activation/Exhaustion Genes, ANOVA comparison for LFR vs. SFR samples
This table lists the results of the ANOVA comparison of 143 activation/exhaustion markers between LFR and SFR cases, using scRNA-seq data.

##### Supplementary Table 10. Activation/Exhaustion Genes, ANOVA comparison for SFR vs. LFR samples in CD4+ T cells only
This table lists the results of the ANOVA comparison of 143 activation/exhaustion markers for CD4+ T cells in the LFR vs. SFR samples.

##### Supplementary Table 11. Clinical characteristics of the 50 AML extension cases
This table cotains clinical information and list the somatic mutation in recurrently mutated AML genes for the 50 extension cases used to validate the immunological findings.

##### Supplementary Table 12. Multivariate analysis for CD4 T cell activator status
This table contains the results for the multivariate analysis of covariates present at diagnosis and potentially associated with CD4 T cell activation.

##### Supplementary Table 13. Neoantigen prediction
This table lists the predicted neoantigen based on somatic mutations detected in the SFR and LFR cases.
